# Mechanistic insights into proteolytic activation of membrane-bound NAC transcription factor SlNACMTF3 in tomato plants under heat stress

**DOI:** 10.64898/2026.10.24.761524

**Authors:** Sushmita Talukdar, Debarati Paul, Shuddhanjali Roy, Amaravadhi Harikishore, Anirban Bhunia, Shubhra Ghosh Dastidar, Pallob Kundu

## Abstract

Stress is perceived at the cellular membrane, where membrane-bound transcription factors (MTFs) are activated, initiating a rapid gene-regulation cascade. We previously demonstrated the multifaceted role of tomato MTF, SlNACMTF3, in regulating expression of biotic and abiotic stress responsive genes. However, the molecular mechanism underlying its activation following stress perception remained unknown. Here, we show that the membrane rhomboid protease SlRP3 recognizes an ISK motif near the transmembrane domain of SlNACMTF3 for cleavage and releasing the transcription factor from the membrane. SlRP3 and SlNACMTF3 co-localize within the same plasma membrane microdomain and interact directly. Molecular dynamics simulations supported the proposed mechanism, in which heat-induced conformational switching of SlRP3, driven by reorientation of the L5 loop, enhances substrate accessibility and proteolytic activity. Furthermore, mutation of key SlRP3 residues involved in substrate interaction impaired SlNACMTF3 activation and reduced the expression of stress-responsive genes. Analysis with SlNACMTF3-overexpressing tomato lines and SlRP3 transient knockdown leaves further revealed that transcription factor activation, stress-responsive gene regulation, and cellular ROS detoxification occur coordinately. Collectively, these findings provide mechanistic insight into stress-triggered SlNACMTF3 activation and identify SlRP3 as a novel membrane-associated stress sensor that activates a major transcriptional regulator to enhance stress preparedness in economically important tomato plants.

**Summary statement:** Heat-dependent conformational change of tomato rhomboid protease 3 facilitates SlNACMTF3 recognition within plasma membrane. The protease-mediated rapid activation leads to stress gene regulation for thermotolerance.

## 1. Introduction

The ability of organisms to respond to environmental cues are fundamentally governed by the regulation of gene expression. In plants, stress-responsive gene expression is tightly controlled by transcription factors (TFs) that perceive internal and external signals either directly or indirectly (Yuan et al., 2019) (Liu et al., 2020). Among plant transcription factors, the NAC (NAM, ATAF1/2, and CUC2) family represents one of the largest and most diverse groups. NAC TFs are involved in a wide range of biological processes, including plant growth, development, and responses to biotic and abiotic stresses. They function as both positive and negative regulators of plant defense against stresses, modulating key processes such as hypersensitive responses and hormone signaling pathways (Amoanimaa-Dede et al., 2022; Kumar et al., 2021; Nuruzzaman et al., 2013; Shao et al., 2015).

A specialized subgroup within this family is the membrane-bound NAC transcription factors (NACMTFs). Unlike conventional TFs, MTFs are initially anchored to cellular membranes in an inactive state. Upon receiving specific stimuli, they are released from the membrane and translocate into the nucleus, where they regulate gene expression. This mechanism enables rapid transcriptional responses, bypassing intermediate processes such as de novo protein synthesis (De Backer et al., 2022). Recent studies highlight the functional significance of NACMTFs. For instance, rice ONAC054 is regulated through membrane release and alternative splicing and is essential for abscisic acid (ABA) signaling during senescence and stress responses (Sakuraba et al., 2020). Similarly, Arabidopsis ANAC089 regulates ABA and redox homeostasis, playing a critical role in seed germination and stress adaptation (Albertos et al., 2021).

In tomato, 13 NACMTFs (SlNACMTFs) have been identified and characterized (Bhattacharjee et al., 2017). Among these, SlNACMTF3 has emerged as a key candidate due to its differential expression across developmental stages and stress conditions. Evidence also suggests that SlNACMTF3 plays a significant role in regulating genes associated with both biotic and abiotic stress responses. The protein comprises 595 amino acids, including an N-terminal NAC domain and a C-terminal transmembrane domain, which enables its membrane localization.

Studies with Arabidopsis suggest regulated intramembrane proteolysis or regulated ubiquitin/proteasome-dependent processing (RIP or RUP) pathway liberates MTFs (De Backer et al., 2022). However, the molecular mechanism of stress-induced activation of RIP or RUP for NACMTF liberation, particularly, in crop plants like tomato, remain poorly understood. Here we investigated the activation or membrane release mechanism of SlNACMTF3 and unveiled RIP process by membrane rhomboid protease (RP) as the regulator of membrane release. Tomato RPs are also stress-responsive (Talukdar et al., 2024) and affect stress physiology of this plant. In this study, a combination of bioinformatic and molecular studies provided insight into the previously unknown details of sensing stress on cellular membrane by RPs and stress signal transmission through SlNACMTF3.

Tomato is an economically important crop and a model system for Solanaceae research (Nejatfard et al., 2021). However, its productivity is severely affected by environmental stresses, especially rising temperatures associated with climate change. Heat stress disrupts reproductive development, reducing fruit set and quality, thereby significantly impacting yield. Understanding stress-responsive regulatory networks is therefore essential for developing resilient crop varieties. Insights gained from this work will contribute to a deeper understanding of plant stress signaling networks and may provide novel targets for genetic improvement.

## 2. Materials and Methods

### 2.1 Plant material, growth conditions and stress exposure

Tomato, *Solanum lycopersicum* L., cultivar Pusa Ruby was used for all studies. Plants were germinated and grown in synthetic soil, Soilrite (Soilrite Mix Tc, Keltech Energies Limited) under moistened conditions. Hoagland nutrient solution or Suphala (NPK=15:15:15) fertilizer was administered to the plants after the emergence of true leaves once a week. The plants were grown in greenhouse under 16hr light/8 hr dark conditions at 25°C, with 70% relative humidity. We have used 30 to 45-day-old plants for each experiment. For heat stress experiments, plants were exposed to 37 °C for 16 hours, and those kept in normal optimum physiological conditions were used as controls. Plants were kept wrapped with plastic covers to prevent excessive water loss during heat stress exposure.

### 2.2 RNA extraction and cDNA synthesis

Total RNA was isolated from 1-1.5-month-old tomato leaf samples of control, mock-inoculated, stressed and various constructs infiltrated tomato leaves using TRIzol reagent (Invitrogen) and following the manufacturer’s protocol. Collected tissue samples were harvested, snap frozen in liquid nitrogen and homogenized using prechilled DEPC-treated double autoclaved mortar-pestle. Quantification of RNA samples was conducted using a nanodrop ND1000 spectrophotometer (Thermo Fisher Scientific, Waltham, MA USA). RNA integrity assessment was conducted by gel electrophoresis of denatured RNA. cDNA synthesis was performed using 5 μg of total RNA, Oligo (dT)18 primers (0.025μg/μl), 1 mM dNTPs and 200 U Thermo Scientific™ RevertAid Reverse Transcriptase along with Thermo Scientific™ RiboLock RNase Inhibitor in 1X RT buffer following the manufacturer’s protocol. cDNA was stored at −20°C.

### 2.3 Real Time PCR analysis

Primers were designed manually, and the properties analyzed using the OligoAnalyzer software. Unique regions were selected for primer designing to ensure specific amplification in RT-PCR analyses. 2.5-5% of the cDNA reaction mix was used as template in 20 μl reaction containing PowerUp™ SYBR™ Green Master Mix and gene-specific primers. Data were normalised based on the expression of the housekeeping EF1α gene transcripts. Relative expression levels of transcripts were determined using the 2^−ΔΔcT^ method.

### 2.4 Agroinfiltration in tomato leaves and onion peels

The CDS of SlRP3 (Solyc03g026080), SlRP7 (Solyc07g042930), and SlRP9 (Solyc09g010880) were cloned into pCAMBIA1304 vector using gene-specific primers for in-planta transient assays (Talukdar et al., 2024). For transient studies, Agrobacterium cells (LBA4404 or GV3101) were transformed with positive clones and allowed to grow until OD_600_ reached 0.6–0.8. For in planta transient assays, three to four-week-old tomato plants were agroinfiltrated with Agrobacterium harbouring desired constructs suspended in infiltration medium (10mM MES-MgCl_2,_ pH 5.6 and 100µM acetosyringone). Onion epidermal cells were infiltrated with transformed Agrobacterium suspension in infiltration liquid containing 41.65 mM D-glucose, 10 mM MES-KOH pH5.6, 10 mM CaCl_2_, 0.05 mM MgCl_2_, 0.011 µM Benzyl amino purine, 100 µM acetosyringone.

### 2.5 GUS reporter assay

Infiltrated leaf samples were crushed in liquid nitrogen in microcentrifuge tubes and suspended in 500 μl of GUS extraction buffer (50mM sodium phosphate buffer pH=7, 10 mM Na_2_EDTA, 10mM β-mercaptoethanol, 0.1% Sarcosyl, 0.1% triton X-100). The tubes were centrifuged at 18,000 g for 20 min at 25°C for clarification. 250 µl of GUS assay buffer (2 mM MUG supplemented to GUS extraction buffer) was mixed with 50 μl of the clear supernatant and the mixture was incubated at 37°C for about an hour. Fluorescence of the solution, due to the production of fluorogenic substrate 4-MU, was measured after exciting at 365 nm and collecting 455 nm emission readings using Varioskan flash multimode plate reader (Thermo scientific, Finland). Protein concentration was measured for each sample using Bradford reagent (Bio-Rad, CA, USA). GUS activity was represented in terms of the amount of MU produced, based on a standard curve, per μg of protein in 1 hour.

### 2.6 Subcellular localization analyses

Full length coding sequences of SlRPs fused with tdtomato at its N-terminus (SlRP-tdTomato), cloned into plant expression vector pCAMBIA1304, where expression is promoted by 35S2 promoter (Talukdar et al., 2024) were used. Localization patterns of SlRPs were compared with the localization of an Arabidopsis plasma membrane marker protein AtPIP2A, which was translationally fused to eGFP at its N-terminus within pCAMBIA1304 vector (AtPIP2A-eGFP) as shown in Supplementary File Figure 1. eGFP-SlNACMTF3 construct as described in one of our previous literature was also utilized in this study (Bhattacharjee et al., 2017). The constructs were co-agroinfiltrated into onion epidermal peels and Leica Sterallis 5 confocal microscope was used for visualisation and imaging 48 hours post-incubation at 28°C. Imaging parameters were kept identical across all experimental groups to ensure linear and accurate comparative quantification. Quantification of eGFP intensity in nucleus was performed using Leica Application Suite X (LAS X) software. Nuclei were segmented to define individual regions of interest (ROIs) based on intensity thresholding of the DAPI channel. These nuclear masks were subsequently mapped onto the corresponding eGFP channel. The Fluorescence Intensity of eGFP within each nuclear ROI was automatically measured, following local background subtraction calculated from cell-free areas.

### 2.7 Bimolecular fluorescence complementation assay (BiFC)

To study the interaction between SlRPs and SlNACMTF3, fragments comprising full-length cDNAs of SlRP3,7 and 9 were cloned in split YFP vector clones (Supplementary File Figure 2), pCAMBIA1304: nEYFP to generate SlRPs-nEYFP constructs. CDS of SlNACMTF3 was cloned in pCAMBIA1304: cEYFP vector to obtain SlNACMTF3-cEYFP. Agrobacterium suspension of LBA4404 strains containing these vector constructs were co-infiltrated in onion epidermal peels and tomato leaves. Agrobacterium cells consisting of various control sets were infiltrated to rule out any false positive results. Leica Sterallis 5 confocal microscope was used for visualization of these peels 48 hours post incubation at 28°C for YFP fluorescence.

### 2.8 Split-ubiquitin yeast two hybrid screening

Coding sequences of SlNACMTF3 and SlRPs were cloned within pNXgate32-3HA and pMetYCgate vectors, respectively. Both putative interactors were co-transformed in ThyAP4 cells were selected onto Leucine and Tryptophan dropout plates. To identify the true interactors, the positive transformants were further screened onto SD/-Leu-Trp-His-Ade plates in the presence of 50 mM 3-aminotriazole (3-AT). The interaction was further substantiated by detecting beta-galactosidase activity using X-gal as substrate by filter lift assay and β-galactosidase activity quantification using 4-methylumbelliferyl β-D-galactopyranoside as another substrate.

### 2.9 Structural modeling of SlRP3, MD Simulations and docking studies

The structure of the Tomato Rhomboid protease SlRP3, with atomic coordinates, is not yet available from any experimental data. Therefore, the structural model of the same was built utilising the homology model method (in the SWISS-MODEL web server), using the homologous protein GlpG from *E. coli* (PDB 2IC8) as a template (Burley et al., 2022; Waterhouse et al., 2018). Unlike GlpG, the Tomato SlRP3 contains a long unstructured loop connecting helices 1 and 2, which was then grafted on the system using PyMOL (v. 3.1.0, Retrieved from http://www.pymol.org/pymol) after modelling this part using AlphaFold(Jumper et al., 2021). Supplementary File Figure 4 A represents the structures of SlRP3. The helices have been marked as H1-H6 and the loops are marked as L1 and L5. The model of the G213A mutant was built by mutating the residue in CHARMM-GUI webserver(Jo et al., 2008).

To embed the protein models in membrane, GBSW implicit membrane model was used, that gives the advantage of carrying out the calculations on protein-membrane systems within a reasonable computing time. The implicit membrane consisted of a 30 Å hydrophobic core slab, with a 5 Å slab on either side to gradually switch the dielectric constant of the medium from membrane to water. The initial orientation of the protein within the membrane was selected arbitrarily yet keeping the axes of the helices more or less aligned with the membrane normal (i.e. vertical to the membrane surface), as shown in Supplementary Figure 4B. After brief energy minimization of the systems, molecular dynamics (MD) simulation of each system was performed separately at two different temperatures, at 298K and 310K using the Langevin dynamics algorithm using CHARMM47b2 software and CHARMM36 force-field (Huang and MacKerell Jr, 2013). Non-bonded interactions were switched to zero between 16 Å and 20 Å. The integration time step was set to 2 fs after applying SHAKE (Ryckaert, Ciccotti and Berendsen 1977) to constraint the vibrations of bonds involving hydrogen atoms. Each trajectory was 100ns long and the data were saved at every 10 ps. The systems are listed in the following Table 1.

**Table 1:** Details of different SlRP3 systems and simulation times.

| Description | System name | Simulation temperature | Simulation length |
| --- | --- | --- | --- |
| Wild-type open conformation at room temperature (residues 33-242) | WON | 298K | 100ns |
| Wild-type open conformation at high temperature (residues 33-242) | WOH | 310K | 100ns |
| G213A mutated open conformation at room temperature (residues 33-242) | MON | 298K | 100ns |
| G213A mutated open conformation at high temperature (residues 33-242) | MOH | 310K | 100ns |

Protein–protein docking methods play a key role in PPI prediction. High-precision docking methods such as HADDOCK (https://rascar.science.uu.nl/haddock2.4/) have been used for docking interaction studies. Peptide stretch (Y565ISKV569) comprising a putative protease cleavage site and one amino acid residue of TM domain of substrate SlNACMTF3 was used for docking purposes with SlRP3. The focus was on SlRP3 since it was highly responsive to thermal stress. The putative interacting residues of SlRP3 (E90, 96 and N197) with peptide stretch of SlNACMTF3 as detected by docking were mutated to amino acid Alanine (E90A, E96A and N197A). The triple mutant protein of SlRP3, named as mSlRP3, was checked for further interaction studies with SlNACMTF3 via Split-Ubiquitin Yeast-2-Hybrid assay and Reporter Assay.

### 2.10 CRISPRi-mediated knockdown of SlRP3

In order to get an insight into the biological significance of SlRP3, transcriptional silencing of SlRP3 was executed utilizing a CRISPR interference (CRISPRi) platform. CRISPRi-based pHSN6I01 vector was used(Alok et al., 2021). Two 20 bp gRNA sequences were selected for each gene. Bioinformatic target prediction tools, CRISPR-P v2.0 and CCtop were used for sgRNA designing. Forward and reverse oligonucleotides were mixed in equal proportions, heated at 95°C for 10 mins, and then kept at room temperature for duplex formation. Annealing created *Bsa*I compatible sites in each end. The duplex was then phosphorylated using T4 PNK and ligated to *Bsa*I digested vector. Clones were confirmed by Sanger’s sequencing and positive clones were transformed into Agrobacterium GV3101:pSOUP competent cells for agroinfiltration infiltration. The SlRP3:CRISPRi systemic construct was delivered to the third and fourth true leaves of tomato seedlings via standard Agrobacterium-mediated infiltration. All subsequent stress treatments, Trypan blue histochemical staining, and physiological quantifications were performed using these SlRP3:CRISPRi plant samples at peak silencing efficiency, which was molecularly validated by qRT-PCR analysis.

### 2.11 Antioxidant enzyme activity assays

Antioxidant enzyme activities were detected by various spectrophotometric assays. Leaves Agroinfiltrated with SlRP3 CRISPRi constructs incubated at 25 °C or subjected to 37 °C heat stress. Tissues were harvested, crushed in liquid Nitrogen and subsequently homogenised in 100 mM potassium phosphate buffer (pH 7.8). The homogenate was centrifuged at 16000 × g for 15-20 min at 4 °C, supernatant was collected for further assays.

#### Superoxide dismutase activity

The method of Steward and Bewley (1980), was used to measure total superoxide dismutase (SOD) activity. One mM riboflavin was added to the reaction mixture containing 100 mM potassium phosphate buffer (pH 7.8), 2 mM EDTA, 9.9 mM methionine, 55 µM nitro-blue tetrazolium (NBT), and 100 µl leaf extract. The mixture containing tubes were illuminated under fluorescent light for 10 min and thereafter absorbance was recorded at 560 nm in a spectrophotometer. One unit of superoxide dismutase activity is defined the amount of enzyme that caused a 50% photochemical reduction of NBT.

#### Catalase activity

Catalase (CAT) activity was determined according to the method described by Aebi (1984). Briefly, the reaction mixture consisted of 100 mM sodium phosphate buffer (pH 7.0), hydrogen peroxide (H₂O₂), and 50 µL of enzyme extract in a final reaction volume appropriate for spectrophotometric measurement. The reaction was initiated by the addition of H₂O₂, and the decomposition of H₂O₂ was monitored by measuring the decrease in absorbance at 240 nm using UV–Vis spectrophotometer. One unit (U) of catalase activity was defined as the amount of enzyme that decomposes 1 μmol of H₂O₂ per minute. Total protein concentration in the sample extract was determined using the Bradford assay with bovine serum albumin (BSA) as the standard. Specific activity was calculated using a molar extinction coefficient of 40 mM⁻¹·cm⁻¹ for H₂O₂ and expressed as units per minute per milligram of protein (U⋅min⁻¹⋅mg⁻¹).

#### Ascorbate peroxidase activity

Ascorbate peroxidase (APX) activity assay was performed according to the protocol described by Nakano and Asada (1981) .100 µl of leaf extract was added to 3 ml of reaction mixture comprising 100 mM potassium phosphate buffer (pH 7.0), 0.5 mM ascorbate and 0.5 mM H_2_O_2_. Enzyme activity was calculated using molar extinction coefficient 2.8 mM^−1^ cm^−1^. One enzyme unit corresponds to amount of enzyme needed to oxidise 1 nmol of ascorbic acid/min/g dry weight.

### 2.12 Electrolyte leakage quantification

Electrolyte leakage was determined as described previously with minor modifications(Chowdhury et al., 2025). The leaf samples prior to conductivity measurement were washed with distilled water to remove dirt from surface. It was then cut into sections and then immersed in tubes containing distilled water of equal volume at room temperature and the first electrical conductivity (ECa) of leaf disks from various samples was noted. The leaf disks thereafter were incubated at 55°C for 25 min, and electrical conductivity (ECb) was measured. At last, test tubes were boiled at 100°C for about 10 min, and the electrical conductivity (ECc) was noted. Electrolyte leakage was determined using the formula: electrolyte leakage (%) = (ECb – ECa)/ECc × 100.

### 2.13 Determination of MDA Content

Malondialdehyde (MDA) content was determined according to the protocol of Ding et al., (2007). 500 mg of tissue was collected from various samples, crushed with liquid nitrogen, and mixed with 0.6% Thiobarbituric acid (TBA) (prepared in 10% trichloroacetic acid). The mixture was then boiled at 100°C for 15 minutes, thereafter, cooled at 4°C and subjected to centrifugation at 10,000g for 20 min. Absorbance of the supernatant at 450 nm, 532 nm, and 600 nm were noted. Extinction coefficient, 155 mM−1 cm−1 was used to estimate the level of lipid peroxidation. Equation used for MDA content estimation was MDA (µM) = 6.45(A532 -A600) −0.56(A450).

### 2.14 Detection of H_2_O_2_ accumulation and content estimation

Hydrogen peroxide detection was performed using 3,3-diaminobenzidine (DAB). The leaf samples were incubated in DAB staining solution (1 mg DAB / ml of 10 mM Sodium Phosphate Buffer, pH=7)), in the dark, overnight at 28°C with constant agitation. The leaves were then dechlorophyllised in a bleaching solution containing ethanol/acetic acid/glycerol (3/1/1, v/v/v) and visualised under white light.

Based on the protocol given by Alexieva et al., (2001) H_2_O_2_ content was measured. Leaf samples were crushed in liquid nitrogen. The powder was homogenized with a solution containing 0.1% w/v trichloroacetic acid (TCA), 1M potassium iodide in 100 mM potassium phosphate buffer (pH 7.0). The samples were incubated at 20-25°C in the dark for 20 min. Absorbance was recorded at 390 nm. Standard curve was prepared with known concentrations of H_2_O_2_ in same solution and represented as mM/gm dry weight.

### 2.15 Proline content estimation

Proline content was determined using a modified method described by Bates et al., (1973). One gm leaf sample was extracted with 5 ml of 75% ethanol(Bates et al., 1973; Chowdhury et al., 2025). 500 µl of the extracted sample was mixed with 1000 µl acidic ninhydrin solution and heated at 90°C, cooled to room temperature and absorbance was measured at 520 nm. Proline content was estimated based on a standard curve prepared with known concentrations of proline in the same solution.

### 2.16 Chlorophylls and total carotenoid content estimation

100-500 mg fresh tissues from various samples were taken and crushed in 4:1 acetone-water mixture, thereafter, centrifuged at 5000 x g for 10 min at 4 °C. Absorbance of the supernatant was recorded at 470 nm, 663 nm and 646 nm. The data was calculated accordingly as mentioned in Chowdhury et al., 2025.

### 2.17 Detection of cell death

The leaf samples were incubated in Trypan blue solution (0.4% in deionized water) for 5–10 min, followed by dechlorophyllisation in bleaching solution as mentioned above. The leaves were then visualised under white light.

### 2.18 Statistics

At least three different plants of the same age group (1-1.5 months) were used for conducting the experiments unless mentioned otherwise. The qRT-PCR analyses usually involved cDNAs prepared from three different RNA samples, and reactions were also set up in triplicate. For conducting protease assays via localization studies, onion epidermal peels were collected from three different biological samples. For each construct reporter assays, transient agroinfiltration was performed in 9 tomato leaves. Similarly, sample size for beta-galactosidase activity was nine too.

Statistical analysis was performed using Microsoft Excel 2020 and GraphPad Prism 8.0. Graphs were prepared using GraphPad Prism 8.0. Results were presented as means ± standard error (SE) and subjected to two-way Anova followed by Tukey’s HSD test or multiple t-test followed by Benjamini-Hochberg False Discovery Rate adjustment, whichever applicable. p≤0.05 was considered significant. In qRT-PCR analysis, the relative expression of genes was calculated using the cT method (2^− ΔΔcT^).

## 3. Results

### 3.1 Rhomboid protease liberates SlNACMTF3 from plasma membrane

SlNACMTF3 remains anchored to the plasma membrane at room temperature or conditions conducive for growth. It is released from the membrane immediately after exposure of tissues to heat stress (Bhattacharjee et al., 2017). Bioinformatic studies have revealed an ISK motif, a putative cleavage site for rhomboid proteases (RPs) (Strisovsky, 2016; Strisovsky et al., 2009), immediate upstream to the TM domain region of SlNACMTF3 (Talukdar et al., 2024). We used eGFP-SlNACMTF3 reporter (Figure 1A, upper panel) to investigate the role of RPs and the ISK motif in heat-induced liberation of SlNACMTF3. Protease inhibitor cocktail or TPCK prevented nuclear migration of SlNACMTF3 in infiltrated onion epidermal peels exposed to heat stress (Figure 1A, lower panel). Proteasomal inhibitors had no effect on the nuclear localization. Furthermore, S567F and K568G mutation completely abolished nuclear localization of eGFP-SlNACMTF3 upon heat stress exposure (Figure 1B). These results indicated that rhomboid proteases are responsible for membrane release of SlNACMTF3.

**Figure 1.**
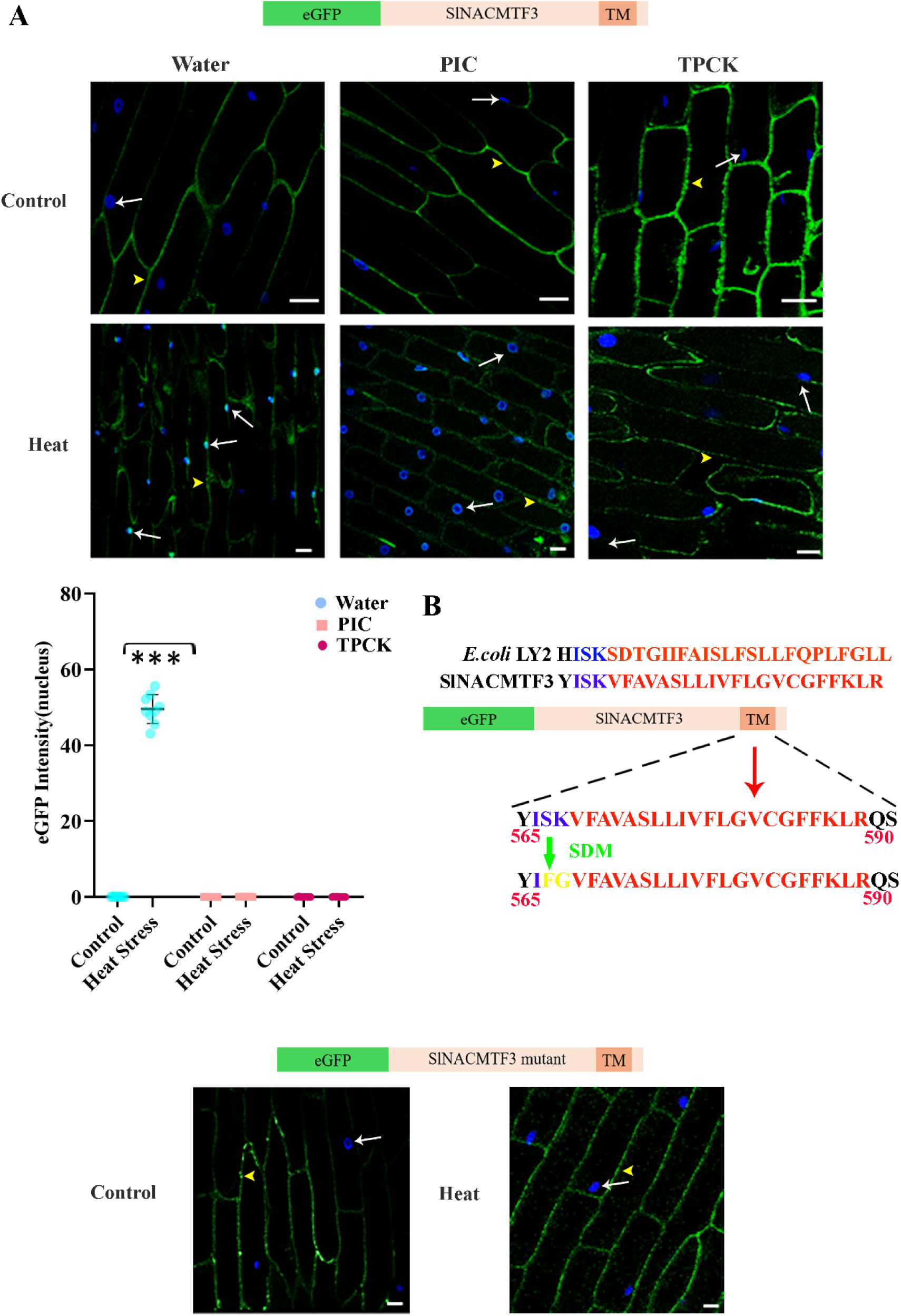
Rhomboid protease-dependent cleavage and nuclear translocation of SlNACMTF3 during heat-stress. (A) Schematic representation of the eGFP–SlNACMTF3 fusion construct used for subcellular localization and proteolytic processing analyses is displayed in the upper panel. SlNACMTF3 contains an N-terminal eGFP tag, the NAC transcription factor domain, and a C-terminal transmembrane (TM) region responsible for membrane association. Middle panel depicts representative confocal micrographs of onion epidermal cells expressing eGFP–SlNACMTF3–TM under control (unstressed, room temperature) and heat-stress conditions, Subcellular localization of GFP (green) in control and heat stressed transformed cells treated with water, protease inhibitor cocktail (PIC) or the serine protease inhibitor TPCK is visible. Nuclei were counterstained with DAPI (blue). White arrows indicate representative nuclei showing GFP accumulation and yellow arrowheads indicate membrane-associated GFP signals. Heat stress promotes clear nuclear accumulation of GFP in mock-treated cells, an effect that is suppressed by both PIC and TPCK treatments. Scale bars = 20 μm. Lower left panel depicts quantification of nuclear GFP fluorescence intensity following control or heat treatment in the presence of water, PIC, or TPCK. Heat stress significantly increased nuclear GFP accumulation in the water-treated samples, whereas this increase was abolished by protease inhibition. Data are presented as mean ± SE from three independent biological replicates: N=3, n>6 nucleus analysed; each dot represents GFP intensity in one nucleus. Statistical significance was determined using Two-way ANOVA followed by Tukey’s multiple-comparison test; ***, P ≤ 0.001 (B) The lower right panel displays Sequence alignment identifying a conserved protease-recognition motif in SlNACMTF3 homologous to the known Rhomboid protease cleavage site of Escherichia coli LY2. The conserved ISK motif (blue) lies within the juxtamembrane region of SlNACMTF3 (residues 565–590); the red font denotes the TM domain vicinal to the protease cleavage site. Site-directed mutagenesis (SDM) was used to substitute the core recognition residues (ISK → IFG, FG in yellow), generating a cleavage-resistant mutant SlNACMTF3 variant. The lower central panel depicts a schematic of the eGFP–SlNACMTF3 mutant construct bearing the SDM-introduced cleavage-site substitution, and representative confocal micrographs of onion epidermal cells expressing this mutant under control and heat-stress conditions. Confocal fluorescence microscopy images of onion epidermal cells show subcellular localisation of the cleavage-site mutant eGFP–SlNACMTF3 construct under control (25°C: normal physiological condition) and heat-stressed conditions(37°C). Mutation of the predicted rhomboid cleavage motif abolishes heat-induced nuclear translocation, with the mutant protein remaining predominantly membrane-associated, confirming that proteolytic cleavage at this site is required for heat-induced nuclear release of SlNACMTF3. GFP fluorescence is shown in green and nuclear staining in DAPI (blue). White arrows indicate GFP in nuclei and yellow arrowheads indicate membrane-associated GFP signals. Scale bars, 20 μm.

### 3.2 Confirmation of SlNACMTF3 activation by rhomboid protease

To confirm the putative role of Rhomboids in the membrane release and activation of SlNACMTF3, we designed a reporter assay (Figure 2A). The reporter construct consisted of core promoter elements of CaMV35S gene and a consensus NAC binding site (NACbs) which would drive GUS expression. We expected SlNACMTF3 would regulate GUS expression from this promoter if activated by SlRPs (Figure 2B) during stress.

**Figure 2.**
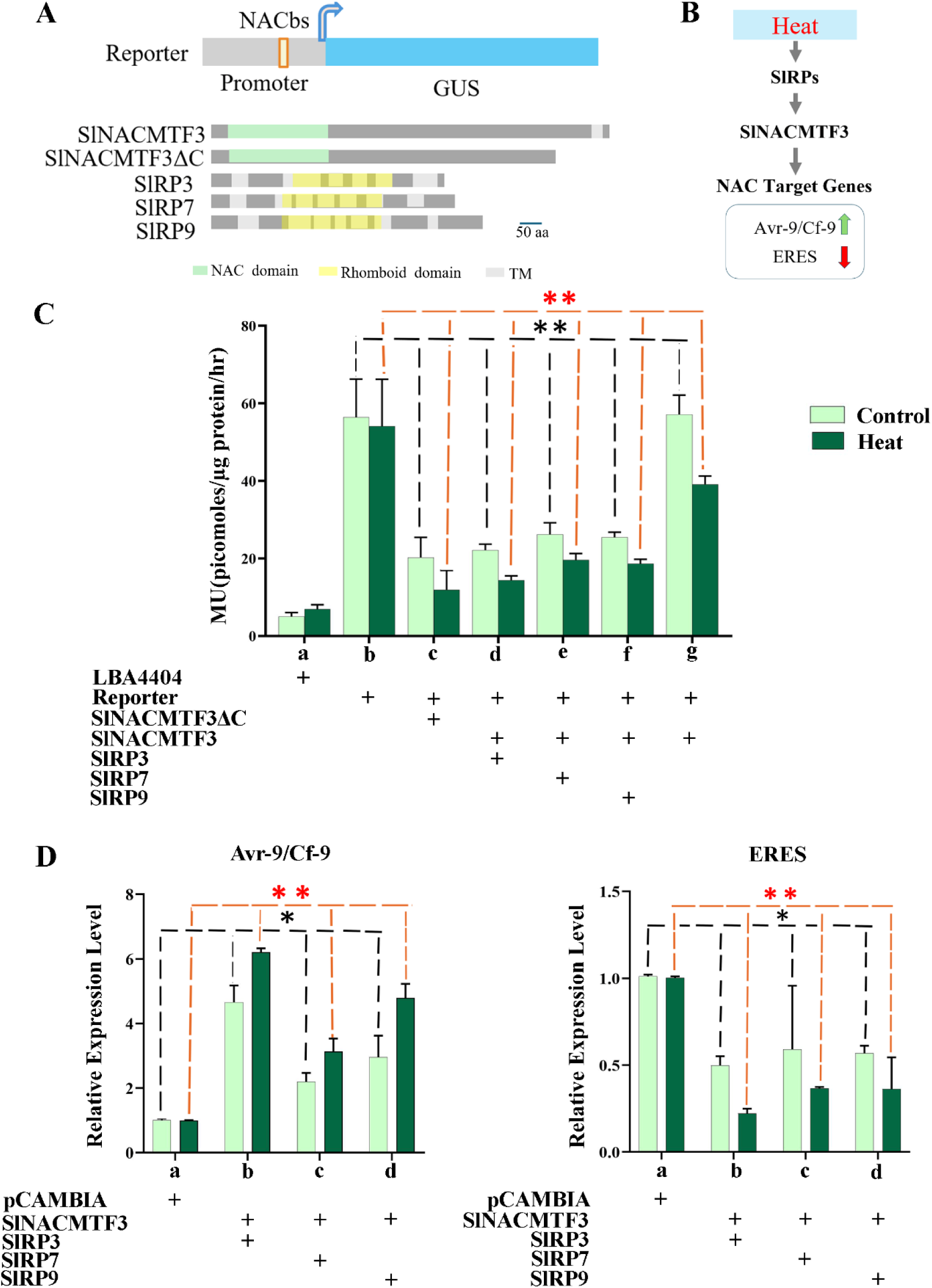
Assessment of SlRPs’ influence on the transcriptional activity of SlNACMTF3. (A) Schematic representation of the constructs used in reporter assays. The reporter construct contains a NAC-binding site (NACbs) in the promoter driving the GUS expression. Domain organization of full-length SlNACMTF3, the truncated lacking the transmembrane (TM) domain, SlNACMTF3ΔC, and tomato rhomboid proteases SlRP3, SlRP7, and SlRP9 is shown. (B) Proposed scheme of events for heat stress-induced activation of SlNACMTF3 and expression of stress-responsive genes Avr9/Cf-9 and ERES. Heat stress activates SlRPs, which proteolytically cleave membrane-tethered SlNACMTF3, releasing the transcription factor for nuclear translocation and subsequent regulation of known NAC-responsive target genes. (C) GUS-activity based reporter assay for assessing SlRP-mediated activation of SlNACMTF3. SlNACMTF3ΔC construct was used here as a positive control. LBA4404 refers to Agrobacterium without any vector. Bars represent mean ± standard error (n=9), * and **symbolizes significant change with p ≤ 0.05 and p<0.001, respectively, as observed by two-way ANOVA followed by Tukey’s multiple comparison test. (D) qRT-PCR data showing differential expression of two endogenous NAC Target genes (Avr-9/Cf-9 and ERES) upon co-infiltration of SlNACMTF3 along with SlRP3 or SlRP7 or SlRP9 in normal physiological conditions (control) and upon heat stress (heat). pCAMBIA1304 refers to agroinfiltration of the empty binary vector pCAMBIA1304. Relative expression of each gene was normalized with the expression level of EF1α. Bars represent mean ± standard error (n = 3), * and ** symbolize significant change p ≤0.05 and p<0.001, respectively, as observed by two-way ANOVA followed by Tukey’s multiple comparison test..

Coinfiltration based assays (Figure 2C) indicated that the extent of promoter repression by SlNACMTF3 in the presence of SlRP was equivalent to the regulation obtained with SlNACMTF3ΔC, which is an activated version of SlNACMTF3. Among the tested SlRPs, the highest repression (3.76 ± 1.40 fold) of the GUS reporter gene was attained upon co-infiltration of SlNACMTF3 with SlRP3, implying this RP has a major role in SlNACMTF3 activation. Sole infiltration of the reporter constructs, without SlNACMTF3, did not display GUS reporter gene repression in both normal physiological conditions and even upon heat stress perception. Heat stress has small but significant influence on activation of SlNACMTF3 or promoter regulation in this assay, perhaps due to the use of overexpression constructs (Figure 2C).

### 3.3 SlRPs influence the expression of endogenous NAC Target genes

We next investigated whether SlRPs could regulate endogenous NAC-target genes. Avr9/Cf9 and Elicitor responsive protein (ERES) are known targets of SlNACMTF3 which gets upregulated and downregulated, respectively, upon SlNACMTF3 activation (Bhattacharjee et al., 2017). qRT-PCR analysis of endogenous transcript level exhibited significant differential regulation of Avr9/Cf9 and ERES upon co-infiltration of SlRPs and SlNACMTF3, further implying occurrence of an interaction among SlRPs and SlNACMTF3. The magnitude of differential regulation of these endogenous NAC target genes increases upon heat stress exposure (Figure 2D). Endogenous transcript levels of Avr-9 got upregulated by 4.6<u>6+</u> 0.52, 2.19±0.26 and 2.96±0.65 folds upon co-infiltration of SlNACMTF3 with SlRP3,7 and 9, respectively, in normal physiological conditions. Elevated expression levels of about 6.20±0.12, 3.13±0.40, and 4.78±0.44 folds were reported upon co-infiltration in heat stressed conditions. Similarly, enhanced downregulation of transcript levels of ERES to 0.22±0.03, 0.36±0.07, 0.36±0.18 folds were enunciated among co-infiltrated samples of SlNACMTF3 and SlRP3, 7 and 9, respectively, upon heat stress as compared to 0.49±0.05, 0.59±0.36 and 0.56±0.04 folds in normal physiological conditions.

### 3.4 SlRPs interact with SlNACMTF3 within plasma membrane

First, we confirmed plasma membrane localization of SlRPs. In onion epidermal peel the red fluorescence signal of SlRP3/7/9-tdtomato overlapped with the green fluorescence signal of the plasma membrane marker AtPIP2a-eGFP (Figure 3A), which suggested that SlRPs resides within the plasma membrane. Upon plasmolysis with 30% (w/v) sucrose, the red fluorescence signal of SlRP3 was separated from the cell wall, which confirmed that SlRP3 was localized into the plasma membrane (Figure 3A). Secondly, the red fluorescence of SlRP3/7/9-tdTomato also overlapped with green fluorescence of eGFP:SlNACMTF3 in plasma membrane, suggesting their close localization as represented in Figure 3B and 3D. The red fluorescence signal of SlRPs didn’t remain diffused throughout the cellular margin and appeared as puncta dots. Over expression of SlRPs also induced nuclear localization of SlNACMTF3, as evident by green fluorescence signal merging with DAPI fluorescence. Highest eGFP intensity in nucleus was obtained upon co-infiltration of SlNACMTF3 with SlRP3 as shown in Figure 3C.

**Figure 3.**
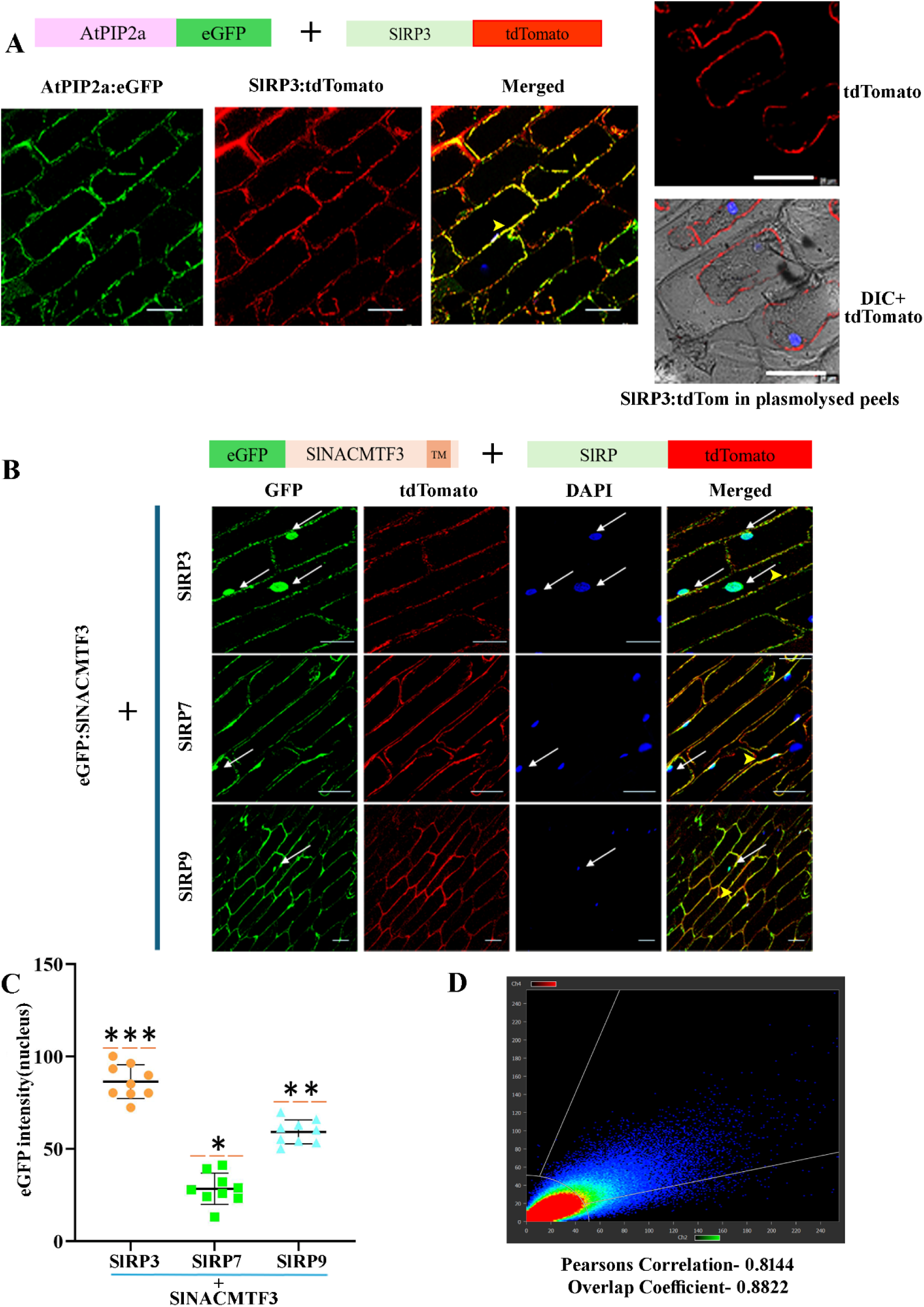
Subcellular localization analysis of SlRPs and SlNACMTF3. (A) Upper panel, outline of the reporter constructs used in subcellular localization analyses in onion epidermal cells. Lower panel shows confocal fluorescence microscopic images for co-localization analysis of AtPIP2a:eGFP, a plasma membrane marker, and SlRP3:tdTomato. The merged images show extensive overlap between the GFP (green) and tdTomato (red) fluorescence in the membrane. Side panel, confocal image of transfected plasmolysed cells depict tdTomato signal remaining associated with the retracted plasma membrane. Scale bars = 20 μm. (B) Representative confocal fluorescence microscopic images for co-localization analysis of eGFP– SlNACMTF3 with tdTomato-tagged tomato rhomboid proteases (SlRP3, SlRP7, and SlRP9) in onion epidermal cells. Diagrams above the image panel indicate constructs used in the study. Nuclei were stained with DAPI (blue). Arrow heads indicate representative regions of co-localization and arrows indicate nuclei. Scale bars = 20 μm. (C and D) Quantification of GFP fluorescence intensity in nuclei shown in lower left panel N = 3, n>3. Each point represents an individual cell, and horizontal bars indicate the mean ± SE. Statistical significance was determined using two-way ANOVA followed by Tukey’s multiple comparison test (**P < 0.01, ***P < 0.001). Lower right panel, representative fluorescence intensity scatter plot used for co-localization analysis between SlRP3 and SlNACMTF3. Pearson’s correlation coefficient (r = 0.8144) and the overlap coefficient (0.8822) indicate a high degree of spatial co-localization between SlRP3 and membrane-bound SlNACMTF3.

To confirm the in vivo interaction of SlRP3,7 or 9 with SlNACMTF3 a bimolecular fluorescence complementation (BiFC) assay was performed using split EYFP protein. SlRP3,7 and 9 were fused to the N-terminus fragment of YFP (SlRP-nEYFP) and SlNACMTF3 was fused to the C-terminus fragment of YFP (SlNACMTF3-cEYFP). The reconstitution of the yellow fluorescence signal was monitored in co-infiltrated tomato and onion epidermal cells. The results confirmed the interaction of SlRP3,7 and 9 with SlNACMTF3 at the membrane vicinity. Absence of YFP signal upon co-infiltration of empty split YFP vector constructs, or SlRP:nEYFP or SlNACMTF3:cEYFP with the empty cEYFP or nEYFP vector constructs (Figure 4A) authenticated BiFC results.

**Figure 4.**
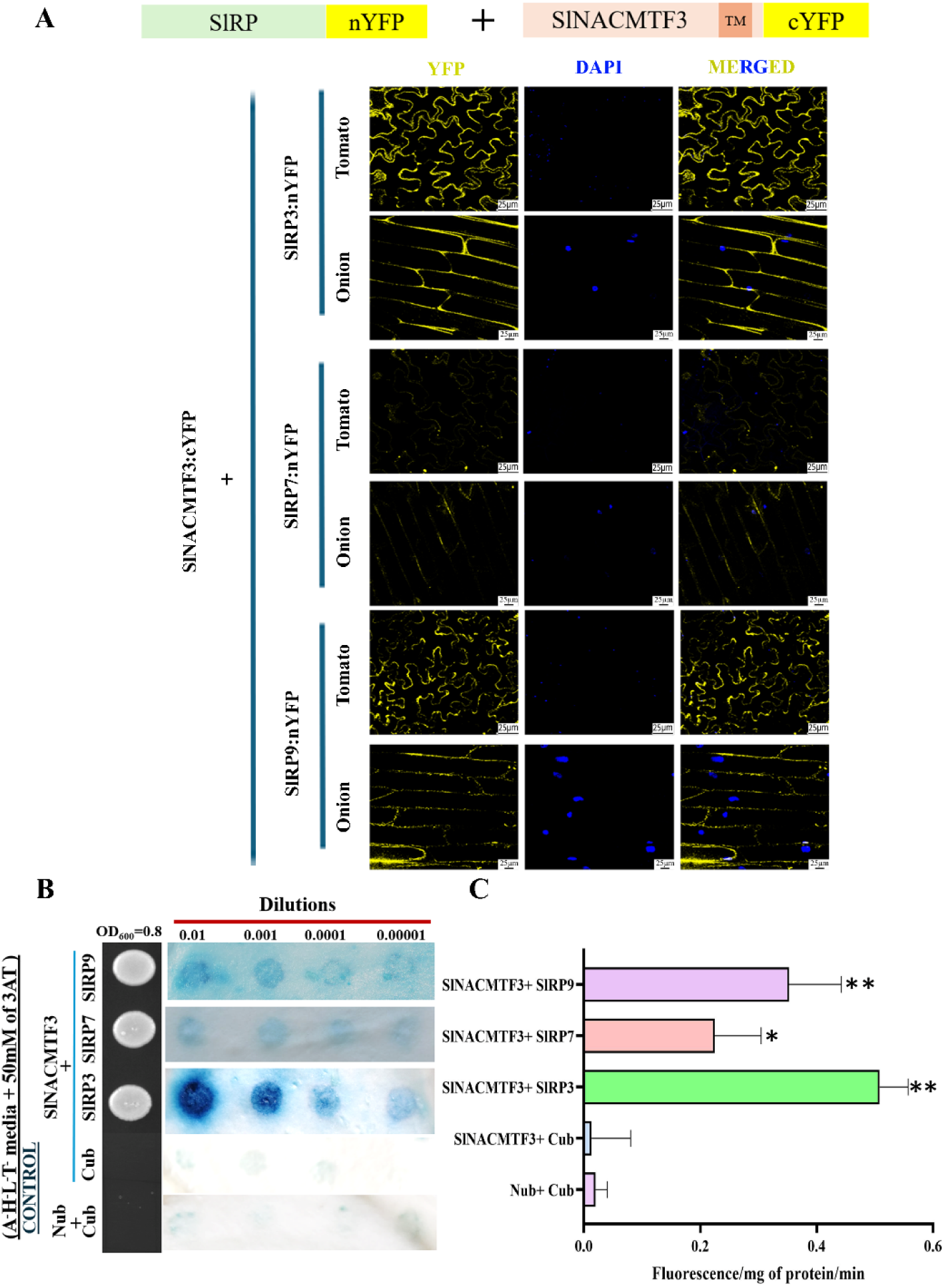
Assays for confirmation of SlRP-SlNACMTF3 direct interaction in vivo. (A) Bimolecular fluorescence complementation (BiFC) analysis demonstrating the interaction between SlNACMTF3 and SlRPs in tomato and onion epidermal cells. Constructs used in the BiFC assays are diagrammatically represented in the upper panel. Reconstituted YFP fluorescence (yellow) indicates protein–protein interaction. Strong YFP fluorescence was observed for the SlRP3–SlNACMTF3 combination, whereas SlRP7 and SlRP9 exhibited comparatively weaker fluorescence. Nuclei were counterstained with DAPI (blue). Merged images are shown in the right panels.Arrowhead represents reconstituted YFP. Scale bars = 25 μm. (B) Yeast split-ubiquitin assay confirming the interaction between membrane-bound SlNACMTF3 and tomato rhomboid proteases. Yeast cells co-expressing SlNACMTF3 with SlRP3, SlRP7, or SlRP9 were serially diluted and grown on selective medium (SD/-Leu/-Trp/-His/-Ade) supplemented with X-Gal and 3-amino-1,2,4-triazole (3-AT). Strong yeast growth and blue color development were observed for the SlNACMTF3–SlRP3 combination, indicating a robust interaction. SlRP7 and SlRP9 displayed weaker reporter activation. Cub and NubG served as negative controls. (C) β-D-galactosidase assay for quantification of interaction strength between two halves of ubiquitin. The SlNACMTF3–SlRP3 interaction exhibited significantly higher reporter activity than the SlNACMTF3–SlRP7 and SlNACMTF3–SlRP9 combinations. Data represent mean ± SE from three independent biological replicates. Statistical significance was determined using one-way ANOVA followed by Tukey’s multiple comparison test (*P < 0.05, **P < 0.01).

### 3.5 SlRP3 is the major SlNACMTF3-interactor protease

Split ubiquitin membrane-based yeast 2-hybrid assay was used to detect SlNACMTF3-SlRPs interactions in membrane vicinity. For the experiment, full-length SlNACMTF3 was fused to mutant N-terminus of split ubiquitin (Nub G), whereas, proteases, SlRP3,7 and 9 were fused to the C-terminus of split ubiquitin. The results indicated that SlRP3, 7, and 9 interacted with SlNACMTF3 as the colonies grew in Ade, His, Trp and Leu quadruple dropout media (Figure 4B). The interaction was further confirmed by beta-galactosidase assay and quantified using 4-methylumbelliferyl β-D-galactopyranoside as substrate which showed highest beta-galactosidase activity with SlNACMTF3 and SlRP3 interactors (Figure 4C).

### 3.6 SlRP3 recognizes the cleavage site in SlNACMTF3

Docking of SlRP3 with peptide stretch (YISKV) comprising putative protease cleavage site and one amino acid residue of TM domain of substrate SlNACMTF3 yielded identification of a few important amino acid residues (E90, E96 and N167) of SlRP3 responsible for the interaction (docking glide score −8.158, dG −66.05 kcal/mol) (Supplementary File Figure 3B). In membrane Y2H assays with E90A, E96A and N197A triple mutant form of SlRP3 colonies did not appear in Ade, His, Trp and Leu quadruple dropout media, thereby suggesting that mutation disrupted SlRP3 interaction with SlNACMTF3 (Supplementary File Figure 3C)

### 3.7 *SlRP3’s conformational change upon thermal stress exposure facilitates cleavage site* recognition and SlNACMTF3 activation

To elucidate the atomistic mechanisms governing SlRP3 substrate recognition under heat stress, MD simulation of SlRP3 was done at different temperatures as detailed in methods section. MD simulation captures atomic-level protein dynamics, providing insights into structure-function relationships. In the structural model of SlRP3, the L5 loop, a short, 7-residue segment that connects helices H5 and H6 (Supplementary File Figure 4A-B) and lies upstream of H225 of the S173-H225 catalytic dyad, which is the putative substrate entry site is similar to bacterial Glpg (Shi et al., 2019). In the initial model, orientation of L5 is oriented away from the dyad (Figure 5A). Visual investigation of the MD simulation trajectories using the VMD software revealed clear differences in the motion of the L5 at different temperatures(Humphrey et al., 1996). In high temperature (WOH), there was not much deviation in the orientation of L5 with respect to their corresponding initial conformation (Figure 5C). In normal temperature (WON), it was interesting to note that, the L5 approached L1 over the course of the simulation, achieving a more ‘closed’ state. That means the L5 remained in an ‘open’ state during temperature stress. To uncover the mechanical driver behind this gating, the residue-specific motion was observed, identifying G213 in the junction between H5 and L5, which possibly acts as a hinge that facilitates local bending. Backbone dihedral angles (Φ, Ψ) of residue 213 were calculated throughout the trajectories (Supplementary File Figure 4D) to understand the molecular mechanism of this ‘gating’. G213, lacking a side chain, exhibited high torsional flexibility, allowing it to sample a region of the Ramachandran space, with dihedral angles clustering between ∼60° to 120° for Φ and ∼0° to 40° for Ψ. Because this part of Ramachandran space is sterically disallowed for standard non-glycine amino acids due to side-chain clashes(Maxwell and Popelier, 2017), this flexibility is likely crucial for coordinating the motions of H5 and L5 during open-to-closed state transitions. To elucidate evolutionarily significance of the ‘gating’, multiple sequence alignment (MSA) was done. Result revealed that G213 is unique in this position (Supplementary File Figure 5A). SlRP7 and SlRP9, homologues of SlRP3 have been reported to function under temperature stress but to a much lesser extents (Talukdar et al., 2024). To further uncover whether the position of G213 at the H5-L5 hinge in the protein structure is unique or not, structures of SlRP7 and S1RP9 were modelled. Overlaid structural representations of the three SlRPs have been presented in Supplementary File Figure 5B. It was clear from the structures that SlRP7 and SlRP9 harbour V195 and A231, respectively, in the corresponding hinge regions. Therefore, it can be hypothesized that the G213 is the principal determinant of this dynamic behaviour. To investigate the role of G213 in RP gating, a G213A mutation was introduced followed by MD simulations under both normal-temperature (MON) and high-temperature (MOH) conditions. Visual investigation of the mutant trajectories showed that the L5 loop preferentially adopted a ‘closed’ conformation under both temperatures. Ala at 213 position introduces a methyl side chain, restricting the accessible torsional space. This residue with high α-helix propensity restricts the backbone to right-handed α-helical dihedral angles in the Ramachandran space. This conformational restriction extends the rigid H5 into the loop boundary, thereby the L5 loop is mechanically drawn inward, stabilizing a closed-loop conformation (Figure 5B-E). The probability distributions of the inter-residue distances of the S173–H225 catalytic dyad were computed across all simulated systems to quantify how this ‘gating’ influences active-site architecture. The inter-residue distances of the S173–H225 catalytic dyad showed distinct profiles across the simulated systems. Probability distribution of the distances revealed that the dyad distance peak was approximately at 5.0 Å for WON. On the contrary, the WOH system displayed a more constricted configuration, with a narrow distribution maxing out at ∼3.0 Å. Peaks of both MON and MOH were shifted towards higher distance ranges (∼ 4.5 Å and ∼4 Å, respectively). Therefore, when the L5 loop remains closed, the dyad distance increases, meaning, the conformational state of the H5–L5 gate directly modulates the organization of the catalytic dyad also. This gating mechanism plausibly determines catalytic competency and substrate accessibility (Figure 5F-J).

**Figure 5.**
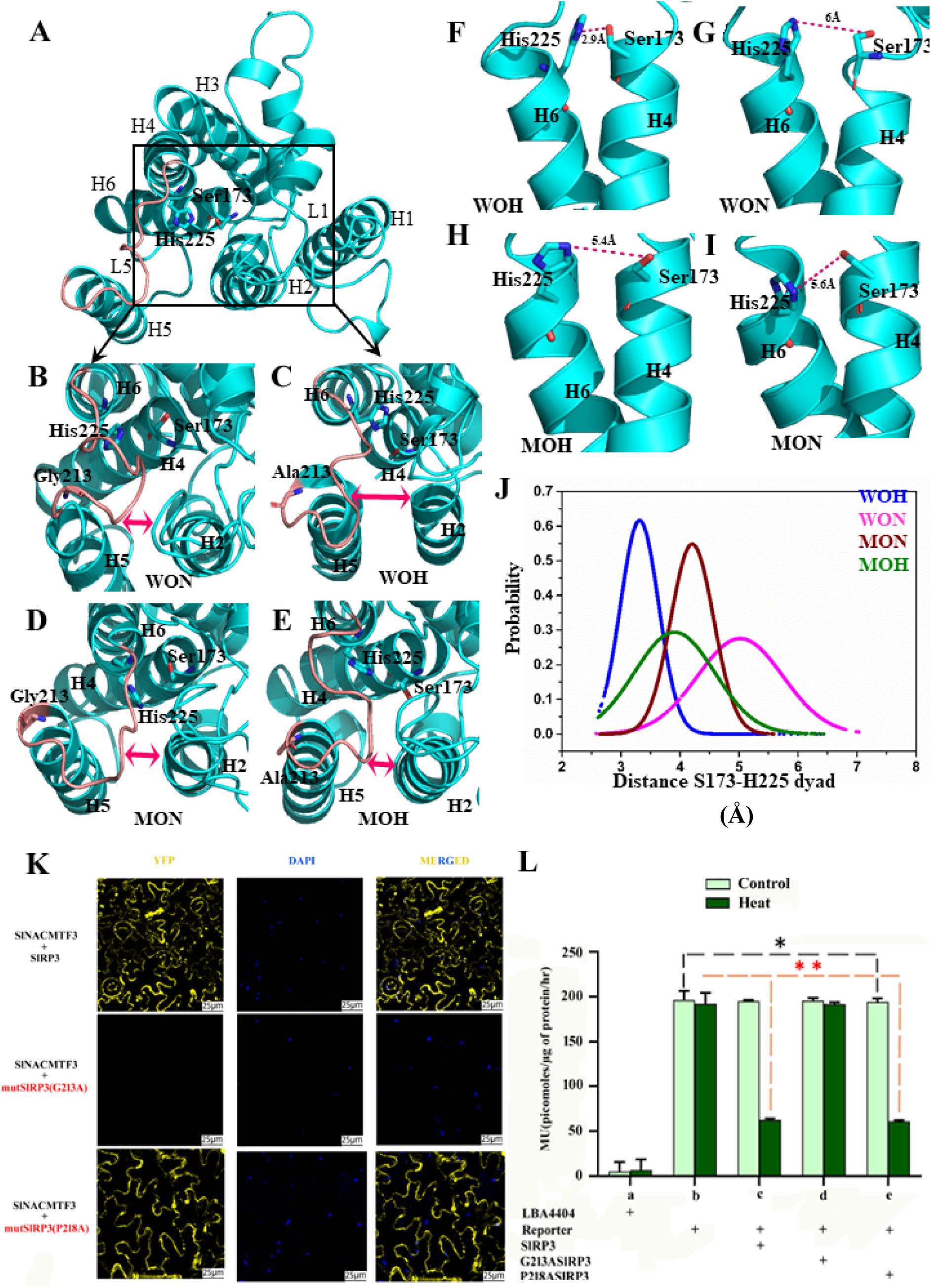
Structural and functional characterization of the catalytic residues and activity of the rhomboid protease. (A) Overall predicted three-dimensional structure showing the transmembrane helices and conserved catalytic residues. (B–E) Structural comparison illustrating the relative positions and interactions of the catalytic residues in the different modelled conditions. (F–I) Close-up views of the conserved catalytic His225–Ser173 dyad under the indicated conditions, with the corresponding inter-residue distances shown. (J) Probability distribution of the Ser173–His225 distance, demonstrating condition-dependent changes in the spatial organization of the catalytic dyad. (K) BiFC analysis of SlNACMTF3 interaction with wild-type and catalytic mutants of SlRP3 in tomato epidermal cells. SlNACMTF3 fused to the C-terminal half of YFP (cYFP) was co-expressed with wild-type SlRP3 (WT SlRP3), the catalytically inactive mutant SlRP3(G213A), or the control mutant SlRP3(P218A) fused to the N-terminal half of YFP (nYFP). Strong YFP fluorescence was detected for the WT SlRP3–SlNACMTF3 and SlRP3(P218A)–SlNACMTF3 combinations, whereas no detectable fluorescence was observed with the catalytic mutant SlRP3(G213A). Nuclei were stained with DAPI (blue). Scale bars = 25 μm. (L) GUS reporter assay for evaluating the effect of wild-type and mutant SlRP3 proteins on SlNACMTF3 activation under control and heat stress conditions. Data represent mean ± SE from three independent biological replicates. LBA4404 indicates transformation with the Agrobacterium without the reporter, served as a negative control. Statistical significance was determined using one-way ANOVA followed by Tukey’s multiple comparison test (*P < 0.05, **P < 0.01).

Taking this cue from MD simulation results, G213A mutation was introduced within SlRP3. Using this G213A mutated form of SlRP3 and wild type SlNACMTF3, BiFC was again conducted at 37°C in tomato epidermal peels to assess their interaction. Absence of reconstituted YFP signal indicated G213A SlRP3 failed to interact with SlNACMTF3 and G213 residue at H5-L5 junction of SlRP3 is crucial for the interaction at higher temperature (Figure 5K). Subsequently, reporter study upon transient co-infiltration of mutant G213A SlRP3 and SlNACMTF3 along with the reporter constructs substantiated the previous observation. Activation of dormant SlNACMTF3 was prevented by this mutated form of SlRP3. An insignificant decrease of 0.98±0.01 fold in GUS reporter activity was observed using G213A SlRP3 compared to 2.26±0.01-fold downregulation with wild type SlRP3 upon heat stress perception (Figure 5L). A P218A mutation was included in the experiment to assess the specificity of the G213A effects. Pro, a residue conserved across the homologs was part of the L5 itself. It is known to restrict the conformations and therefore, the P218A mutation can contribute to the increased flexibility of the L5. However, the P218A substitution did not alter protein function. The P218A mutation construct coinfiltrated with SlNACMTF3 could successfully reduce GUS reporter gene expression by about 2.16± 1.70 fold. Hence, the P218A substitution did not alter protein function (Figure 5L). The lack of functional impact of P218A supports the interpretation that the observed effects are not due to nonspecific mutational perturbations but are instead linked to the unique mechanical role of G213.

### 3.8 Coordinated stress induced expression of proteases and NAC target genes in wildtype and SlNACMTF3 overexpressing plants

Since the synchronization of SlRP3 and SlNACMTF3 activity leads to expression of stress response related genes we investigated whether their stress-dependent expression patterns also match. We monitored expression levels of SlRP3/7/9, SlNACMTF3, Avr/Cf9 and ERES, at same time points post stress application, in wild-type and SlNACMTF3 overexpressing transgenic tomato lines in conditions conducive for growth and upon subjecting the plants to heat stress (37°C/16hrs).

SlRP3, 7 and 9 responded to thermal stress stimulus by exhibiting a gradual increase in their expression levels with the duration of the heat stress in wild-type plants. The magnitude of expression enhancement was higher in SlNACMTF3-overexpressed transgenic plants exposed to heat stress (Figure 6). Changes in the expression of NAC-target genes was not discernible in wild-type plants, however, differential regulation of NAC target genes was prominent in SlNACMTF3-overexpressed transgenic plants receiving thermal stress, while the expression level of SlNACMTF3 slightly enhanced in the transgenic plants. Hence, stress-induced expression of endogenous SlRPs and tested NAC target genes seems to follow a coordinated pattern (Figure 6). The induction of NAC target genes occurred at early time points, which matched with the SlRP activation for releasing SlNACMTF3.

**Figure 6.**
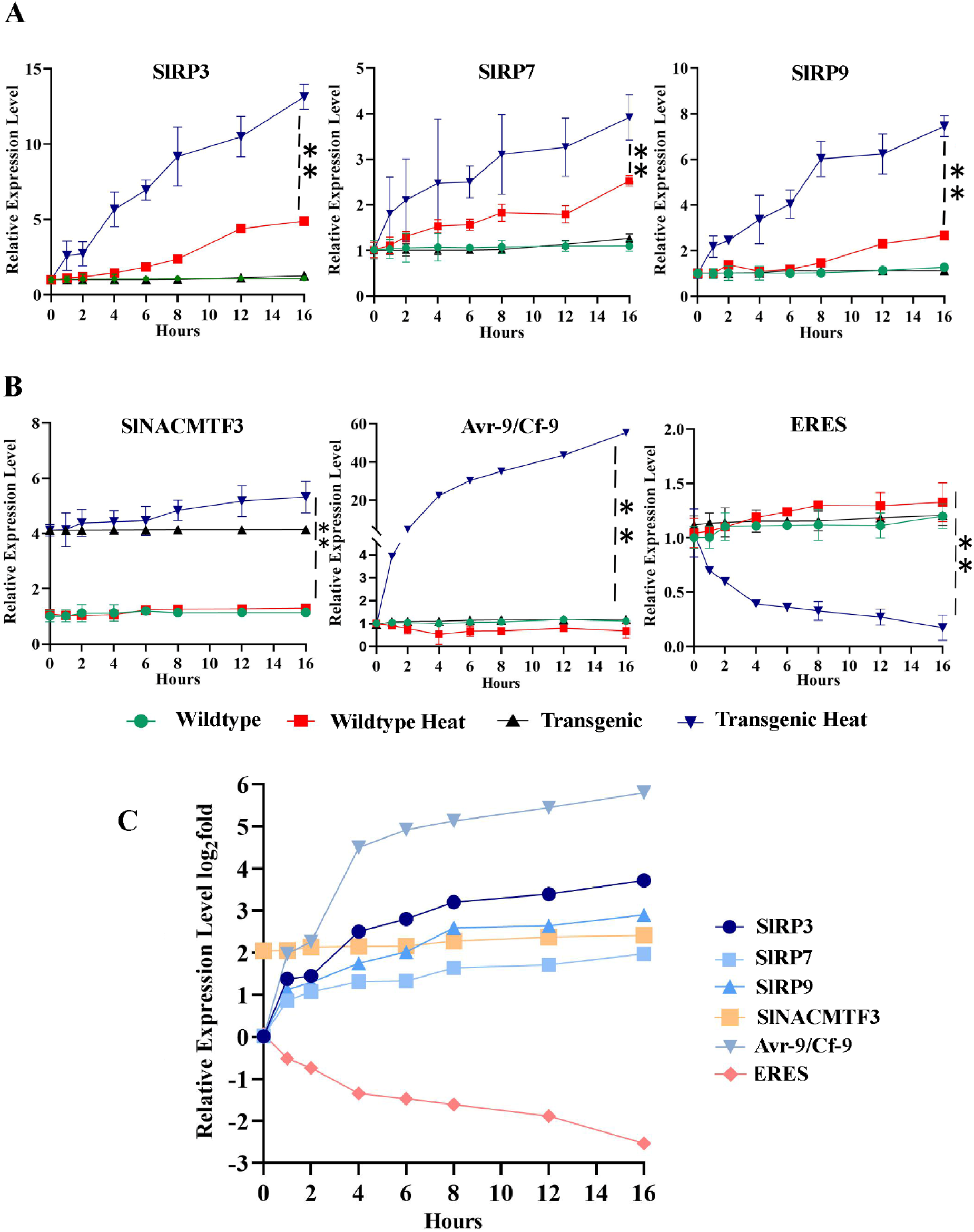
Relative expression level of SlRPs, SlNACMTF3 and NAC target genes in early time points after heat stress exposure of wildtype and SlNACMTF3 overexpressed transgenic tomato plants. (A) and (B) qRT-PCR data depicting transcript expression levels of endogenous SlRPs, SlNACMTF3, Avr9/Cf9 and ERES in wildtype and SlNACMTF3 overexpressing lines at various time points after heat stress exposure. Relative expression of each gene was normalized with the expression level of EF1α. Heat stress induces markedly stronger and more sustained upregulation of SlRP3, SlRP7, SlRP9, SlNACMTF3, and Avr-9/Cf-9 in transgenic plants relative to wild-type, whereas ERES expression is progressively downregulated in heat-stressed transgenic plants but remains largely unchanged in the other three groups. Data are presented as mean ± SE (n = 3 biological replicates); **P < 0.005 (transgenic heat-stressed vs. wild-type heat-stressed). Statistical significance between heat-treated wild-type and SINACMTF3 transgenic plants was determined using a two-tailed paired Student’s t-test, with P-values adjusted for multiple comparisons using the Holm–Šídák method. (C) Comparison of dynamics of expression of SlRPs, SlNACMTF3 and endogenous NAC target genes in SlNACMTF3 overexpressed transgenic lines upon heat stress exposure. Summary plot of relative expression levels (log₂-transformed) relative expression values summarizing the temporal transcriptional behavior of the six genes analyzed, SlRP3, SlRP7, SlRP9, SlNACMTF3, Avr-9/Cf-9, and ERES over the 16 h heat-stress time course in transgenic plants. Together, the expression profiles reveal coordinated and gene-specific transcriptional reprogramming associated with SINACMTF3 during heat stress. Data represent mean ± SE (n = 3).

### 3.9 Transient knockdown of SlRP3 in tomato leaves renders them susceptible to thermal stress

We targeted the promoter region of SlRP3 with dCas9-KRAB for downregulation. qRT-PCR analysis confirmed almost 85% downregulation of SlRP3 transcript level using both the guideRNAs, gRNA1 and 2 as shown in Supplementary File Figure 6. The SlRP3: CRISPRi plant samples showed increased curling and wilting compared to mock-infiltrated wild-type heat-stressed leaves (Figure 7A). Trypan blue and DAB staining confirmed an increase in cell death with concomitant enhanced H_2_O_2_ accumulation in SlRP3:CRISPRi leaves (Figure 7B-C). While SlRP3: CRISPRi renders hypersensitive to heat stress in tomato plants, transient agrobacterium mediated overexpression of the SlNACMTF3ΔC construct effectively rescued this phenotype even in SlRP3: CRISPRi, significantly suppressing heat-induced cellular death and attenuating H_2_O_2_ accumulation (Figure 7A-C). These observations were further corroborated by different biochemical assays which displayed significantly higher H_2_O_2_ accumulation in SlRP3:CRISPRi leaf samples upon thermal stress as compared to mock-infiltrated stressed samples (Figure 7D). ROS usually causes impaired damage to macromolecules that are critical in maintaining cell integrity owing to their highly reactive nature. Heat stress markedly enhanced antioxidant capacity in mock-infiltrated plants, as reflected by a coordinated increase in catalase (CAT), superoxide dismutase (SOD), and ascorbate peroxidase (APX) activities. In contrast, SlRP3:CRISPRi samples severely compromised this antioxidant response, with all three enzyme activities remaining significantly lower than those of heat-stressed mock controls (Supplementary File Figure 7). These observations indicate an essential role of SlRP3 in activating enzymatic ROS-scavenging pathways during thermal stress.

**Figure 7.**
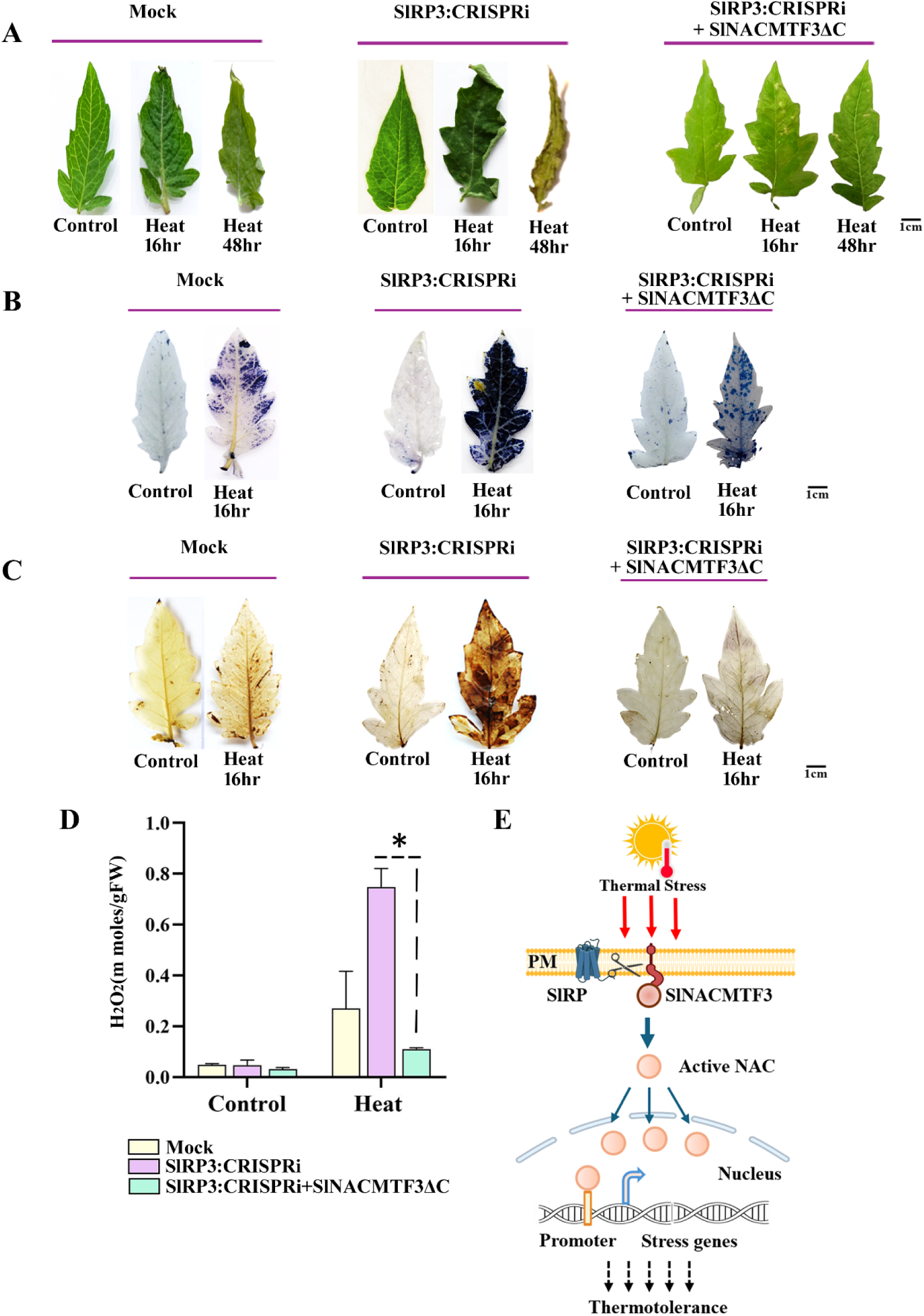
Testing the effect of SlRP3 knockdown on heat tolerance property of tomato. (A) Representative leaf phenotypes of mock and SlRP3: CRISPRi plants, with or without SlNACMTF3ΔC complementation, under control and heat-stress conditions (16 and 48 h). (B) Trypan blue staining (left) was used to detect dead cells under control and heat stress in the indicated genetic backgrounds. (C) Visualization of heat-induced H₂O₂ accumulation in mock, SlRP3:CRISPRi and SlRP3:CRISPRi + SlNACMTF3ΔC plants. (D) Quantification of H₂O₂ levels demonstrating enhanced ROS accumulation in SlRP3:CRISPRi plants following heat stress and its reduction upon expression of the constitutively active SlNACMTF3ΔC. \**Asterisks indicate statistically significant differences (P < 0.05).* (E) Proposed model depicting the role of SlRP3 in heat stress-induced proteolytic release of membrane-bound SlNACMTF3. Heat stress activates SlRP3-mediated cleavage of SlNACMTF3 at the plasma membrane, releasing the transcriptionally active NAC domain, which translocates to the nucleus and regulates stress-responsive genes to promote thermotolerance.

Consistent with impaired stress acclimation, SlRP3:CRISPR silenced samples accumulated substantially less proline under heat stress, whereas mock-infiltrated plants exhibited pronounced proline induction relative to non-stressed controls. Reduced proline levels in knockdown plants likely weakened osmotic adjustment and ROS detoxification capacity. Concomitantly, lipid peroxidation, as measured by malondialdehyde (MDA) content, was strongly elevated in SlRP3:CRISPRi leaves upon heat exposure, while mock-infiltrated leaves maintained significantly lower MDA levels, indicating reduced membrane damage. Electrolyte leakage assays further corroborated these findings, revealing a pronounced loss of membrane integrity in SlRP3:CRISPRi leaves under heat stress compared to mock-infiltrated stressed samples (Supplementary File Figure 7). In agreement with increased cellular damage, photosynthetic pigment content was markedly reduced in SlRP3:CRISPRileaves following heat stress, including chlorophyll a, chlorophyll b, and carotenoids, whereas mock-infiltrated leaves retained comparatively higher pigment levels (Supplementary File Figure 7).

At the transcriptional level, heat-induced upregulation of key antioxidant genes—including SlCAT1, SlCAT2, SlCAT3, SlCSD1, SlAPX1, and SlAPX2—observed in mock-infiltrated leaves was significantly attenuated in SlRP3:CRISPRi samples Notably, SlCSD2 showed an opposite trend, displaying enhanced expression in SlRP3:CRISPRi samples relative to mock controls. These data suggest selective rewiring of antioxidant gene regulation in the absence of SlRP3.

Moreover, SlRP3:CRISPRi resulted in broad suppression of heat shock transcription factors (SlHSFA1a, SlHSFA2, SlHSFA3, SlHSFA6b, and SlHSFB1) under heat stress (Supplementary File Figure 6). Correspondingly, the induction of downstream heat shock proteins (SlHSP20, SlHSP70, and SlHSP90) was only modest in knockdown plants, indicating an impaired heat shock response.

Collectively, these results demonstrate that SlRP3 is a critical regulator of heat stress tolerance in tomato, functioning upstream of antioxidant defenses, Osmoprotectant accumulation, membrane stability, and heat shock signaling pathways. Overall, the data represented in Supplementary table 1 indicates that reduction of levels of SlRP3 in tomato plants renders the plant ‘heat susceptible’(Figure 7E).

## 3. Discussion

The plasma membrane serves as the primary interface between plant cells and their environment. Signal perception by membrane-associated receptors initiates complex signaling cascades involving protein phosphorylation, calcium fluxes, reactive oxygen species (ROS) generation, and secondary messengers, ultimately leading to physiological responses such as growth, development, defense, stress tolerance, nutrient acquisition, and environmental adaptation. Signal integration is often accompanied by large-scale transcriptional reprogramming mediated by diverse transcription factors. Membrane-tethered transcription factors (MTFs) add an additional layer of regulatory complexity because they can directly translocate to the nucleus following activation and rapidly regulate target gene expression. Although regulated intramembrane proteolysis (RIP) and regulated ubiquitin/proteasome-dependent processing (RUP) are recognized mechanisms controlling MTF activation (Chi et al., 2017; Tian et al., 2017), limited understanding of plasma membrane-associated RIP and RUP mechanisms during stress has hindered our comprehension of MTF-mediated signaling networks.

In the present study, we demonstrated that (i) treatment with TPCK and mutation of the putative rhomboid protease (RP) cleavage motif (ISK) abolished nuclear translocation of SlNACMTF3; (ii) the transcriptional activity of SlNACMTF3 was dependent on SlRP function (Talukdar et al., 2024); (iii) SlRPs and SlNACMTF3 co-localize within the same plasma membrane microenvironment; and (iv) SlRPs physically interact with SlNACMTF3 in both yeast and plant cells. Collectively, these findings provide compelling evidence that rhomboid proteases mediate cleavage of the transmembrane domain of SlNACMTF3 during thermal stress, thereby regulating its transcriptional activity. Among the members analyzed, SlRP3 consistently emerged as the predominant interactor. These findings agree with previous observations in *Arabidopsis thaliana*, where the endoplasmic reticulum-localized transcription factor ANAC013 was shown to be activated through cleavage by RHOMBOID-LIKE 2 under stress conditions (Eysholdt-Derzsó et al., 2023).

Rhomboid proteases catalyze intramembrane proteolysis of substrates containing a single transmembrane domain, a characteristic feature of SlNACMTF3. Because purification and biochemical characterization of membrane proteins remain technically challenging, we employed molecular dynamics simulations to gain mechanistic insight into the stress-dependent association between SlRP3 and SlNACMTF3 within the membrane environment and the subsequent cleavage of the SlNACMTF3 transmembrane domain. The precise pathway through which substrates gain access to rhomboid proteases remains incompletely understood. Studies on the bacterial rhomboid protease GlpG from *Escherichia coli* suggest that transmembrane helix 5 (TMH5) functions as a lateral gate that permits substrate entry into the catalytic site (Baker et al., 2007; Ha et al., 2013). Furthermore, Wang and colleagues reported that the free-energy difference between the open and closed conformations of Loop 5 (L5) in GlpG is relatively small, facilitating rapid conformational transitions (Wang and Ha, 2007; Wang et al., 2016).

In rhomboid proteases, L5 constitutes a short loop connecting helices H5 and H6 and is located opposite to Loop 1 (L1), adjacent to the catalytic dyad formed by S173 and H225. Therefore, the orientation of L5 appears to be crucial for substrate access. In the closed conformation, L5 is directed toward L1 and lies in close proximity to the catalytic dyad, whereas in the open conformation it is oriented away from the active site, creating a pathway for substrate entry (Ben-Shem et al., 2007; Lemieux et al., 2007; Wu et al., 2006). Structural studies of *E. coli* GlpG have similarly revealed dynamic transitions between open and closed conformations, with coordinated movements of L5 and Transmembrane H5 regulating access to the catalytic center. The temperature-induced conformational changes observed in SlRP3, together with the involvement of residues within the L5 region, are consistent with these established models and suggest that L5 governs substrate accessibility during thermal stress, ultimately facilitating activation of SlNACMTF3.

Our simulations further revealed a pronounced temperature-dependent shift in L5 dynamics. At lower temperatures, the loop predominantly sampled a closed conformation, whereas elevated temperatures promoted a dynamic equilibrium between open and closed states. Residue-level analysis identified G213 as a critical determinant of this temperature-sensitive behavior. Multiple sequence alignment demonstrated that Glycine at this position is highly conserved, suggesting evolutionary selection for local flexibility within an otherwise structurally constrained region. The intrinsic torsional freedom conferred by G213 enables L5 to undergo conformational transitions while maintaining structural integrity. Increased thermal energy further enhances backbone fluctuations, thereby facilitating this dynamic equilibrium. Consistent with this interpretation, temperature-dependent changes in the distances between L1-L5 residue pairs and between residues comprising the catalytic dyad correlated strongly with L5 movement. Together, these findings indicate that G213 plays a key mechanistic role in maintaining the conformational plasticity required for L5 gating. Such flexibility is likely essential for balancing substrate accessibility with optimal positioning of catalytic residues during proteolysis.

During stress, interaction between SlRPs and SlNACMTF3 activates the latter and regulates downstream effector genes, including *ERES* and *Avr3/Cf9*. Regulatory feedback loops are a common feature of stress-signaling pathways. Our detailed expression analyses suggest that (i) activation of SlNACMTF3 represents an early event in the stress-response cascade, (ii) SlNACMTF3 likely influences the expression of SlRP genes, and (iii) the expression patterns of SlRP and SlNACMTF3-associated genes are strongly correlated. Taken together, these findings support a signaling model in which stress-induced activation of SlRPs triggers SlNACMTF3-mediated transcriptional responses, while SlNACMTF3 subsequently amplifies the signaling cascade by promoting SlRP expression. The presence of NAC-binding sites within the promoters of *SlRP3*, *SlRP7*, and *SlRP9* further supports this regulatory feedback mechanism(Talukdar et al., 2024).

A central question arising from this study concerns the physiological significance of the SlRP-SlNACMTF3 signaling axis. Given their strong stress responsiveness, we investigated their role in thermotolerance. CRISPRi-mediated knockdown of *SlRP3* rendered tomato plants significantly more susceptible to heat stress, whereas expression of active SlNACMTF3 substantially restored heat tolerance. These findings indicate that the SlRP3-SlNACMTF3 module is an essential component of the heat-stress response. Efficient scavenging of ROS is crucial for cellular survival under stress conditions. Consistent with this concept, elevated ROS accumulation in *SlRP3* knockdown plants was markedly reduced following SlNACMTF3 complementation. Thus, activation of the SlRP3-SlNACMTF3 pathway appears to regulate key stress-responsive genes that collectively alleviate oxidative damage. In contrast, loss of SlRP3 function results in excessive H₂O₂ accumulation that exceeds the antioxidant capacity of the cell, ultimately leading to oxidative stress.

Exposure to heat stress compromises membrane integrity by altering membrane fluidity, destabilizing lipid bilayers, and accelerating lipid peroxidation, leading to increased malondialdehyde (MDA) accumulation. Loss of membrane stability consequently results in increased electrolyte leakage. Reduced MDA content and lower electrolyte leakage are widely recognized indicators of enhanced thermotolerance and improved ROS management in plants. Similarly, proline accumulation contributes significantly to heat adaptation by functioning as an osmoprotectant and ROS scavenger (Hayat et al., 2012). Antioxidant enzymes such as superoxide dismutase (SOD), catalase (CAT), and ascorbate peroxidase (APX) form the first line of defense against oxidative damage and are essential for maintaining redox homeostasis during heat stress (Mishra et al., 2023). In agreement with previous reports in heat-tolerant genotypes of *Brassica*, lentil, rice, and wheat, improved stress tolerance correlates strongly with sustained antioxidant activity and maintenance of cellular redox buffers. These protective mechanisms restrict the accumulation of hydrogen peroxide and lipid peroxidation products, resulting in superior preservation of chlorophyll content and membrane integrity (Rao et al., 2025).

The involvement of rhomboid proteases in ROS-mediated stress signaling has previously been demonstrated through the release of ANAC017 from endoplasmic reticulum membranes in response to H₂O₂ accumulation (Ng et al., 2013). Previous studies have also established that excessive H₂O₂ can carbonylate proteins and oxidize thiol groups in antioxidant enzymes, thereby reducing their catalytic efficiency (Siemieniuk et al., 2025). Under prolonged stress conditions, antioxidant defense systems can become overwhelmed or functionally impaired, causing CAT and APX activities to become insufficient for maintaining ROS homeostasis (Zandi and Schnug, 2022). Once ROS generation exceeds detoxification capacity, oxidative damage accumulates in membranes, proteins, and nucleic acids and may ultimately trigger programmed cell death (Dmitrieva et al., 2024). Therefore, the inability of *SlRP3* knockdown plants to sustain antioxidant enzyme activities under heat stress likely reflects a disruption of redox regulation resulting from impaired stress signaling. Collectively, these observations indicate that loss of SlRP3 compromises activation of downstream stress-response machinery required for effective heat adaptation.

Heat Shock Transcription Factors (HSFs) constitute another major regulatory layer governing plant thermotolerance by orchestrating the heat shock response (HSR) (Bakery et al., 2024). In tomato, thermotolerance is regulated by a complex network of 21 HSFs, among which HSFA1, HSFA2, and HSFB1 serve as key regulators of heat-responsive gene expression (Singh et al., 2022). HSFA6b functions as an important upstream activator and facilitates interactions among multiple HSF family members. In wheat, HSFA6b has been identified as a central hub within the HSF regulatory network (Huang et al., 2016), while studies in *Arabidopsis* have shown that HSFA6b interacts with HSFA2 and HSFA3. Furthermore, HSFB1-mediated regulation of heat-responsive genes is dependent upon interactions involving HSFA1a, HSFA2, HSFA6b, and HSFA9. HSFs transcriptionally activate Heat Shock Proteins (HSPs), molecular chaperones that prevent protein aggregation, maintain proteostasis, and promote recovery from heat-induced cellular damage. Increased accumulation of HSP70, HSP90, and HSP101 is strongly associated with enhanced thermotolerance (Vierling, 1991). Interactions between HSP70/HSP90 and HSFs further strengthen protective signaling pathways and facilitate activation of downstream heat-responsive genes (Hahn et al., 2011). HSP70 and HSP90 have also been implicated in maintaining photosynthetic function and activating thermotolerance-associated processes under adverse environmental conditions (Donnelly et al., 2013; Fernández-Crespo et al., 2022; Park and Seo, 2015). Likewise, constitutive expression of HSP101 substantially improves survival under extreme temperature stress in both *Arabidopsis* and maize. Therefore, the marked reduction in transcript abundance of *HSFA1a*, *HSFA2*, *HSFA3*, *HSFA6b*, *HSFB1*, *HSF20*, *HSP70*, and *HSP90* observed in *SlRP3* knockdown plants strongly suggests that SlRP3 functions upstream of the HSF-HSP regulatory network and contributes to thermotolerance through modulation of heat stress signaling pathways.

Taken together, our findings establish SlRP3 as a critical upstream signaling component that activates the membrane-bound transcription factor SlNACMTF3 through regulated intramembrane proteolysis. This activation initiates transcriptional programs that reinforce antioxidant defenses, maintain redox homeostasis, preserve membrane integrity, and promote heat-stress adaptation. Consistent with the conserved role of rhomboid proteases in releasing membrane-tethered signaling factors (Grieve et al., 2021; Eysholdt-Derzsó et al., 2023; De Clercq et al., 2013), the SlRP3-SlNACMTF3 module appears to function as a stress-responsive signaling switch in tomato (John et al., 2024). The elevated MDA levels, increased electrolyte leakage and H₂O₂ accumulation, reduced proline content, diminished antioxidant enzyme activities, and decreased expression of HSFs and HSPs observed in *SlRP3* knockdown plants collectively demonstrate a previously unrecognized role of SlRP3 as both a stress sensor and a positive regulator of thermotolerance. Future studies aimed at identifying the direct downstream targets of SlNACMTF3 and elucidating its integration with established heat-stress regulatory networks will further clarify the molecular mechanisms by which the SlRP3-SlNACMTF3 signaling axis confers thermal resilience in tomato.

## Author Contribution

ST, PK conceptualized; ST, SR performed major experiments; DP, SGD conducted MD simulation studies; AH, AB conducted docking analysis, ST, DP, SR, PK wrote the manuscript, all authors have seen the manuscript and edited. PK is responsible for overall management of the project.

## Acknowledgement

Bose Institute Central Instrument Facility, DST, DBT and ANRF GoI, CSIR, GoI

