## Supplementary figures and table 1 for "Mechanistic insights into proteolytic activation of membrane-bound NAC transcription factor SlNACMTF3 in tomato plants under heat stress"

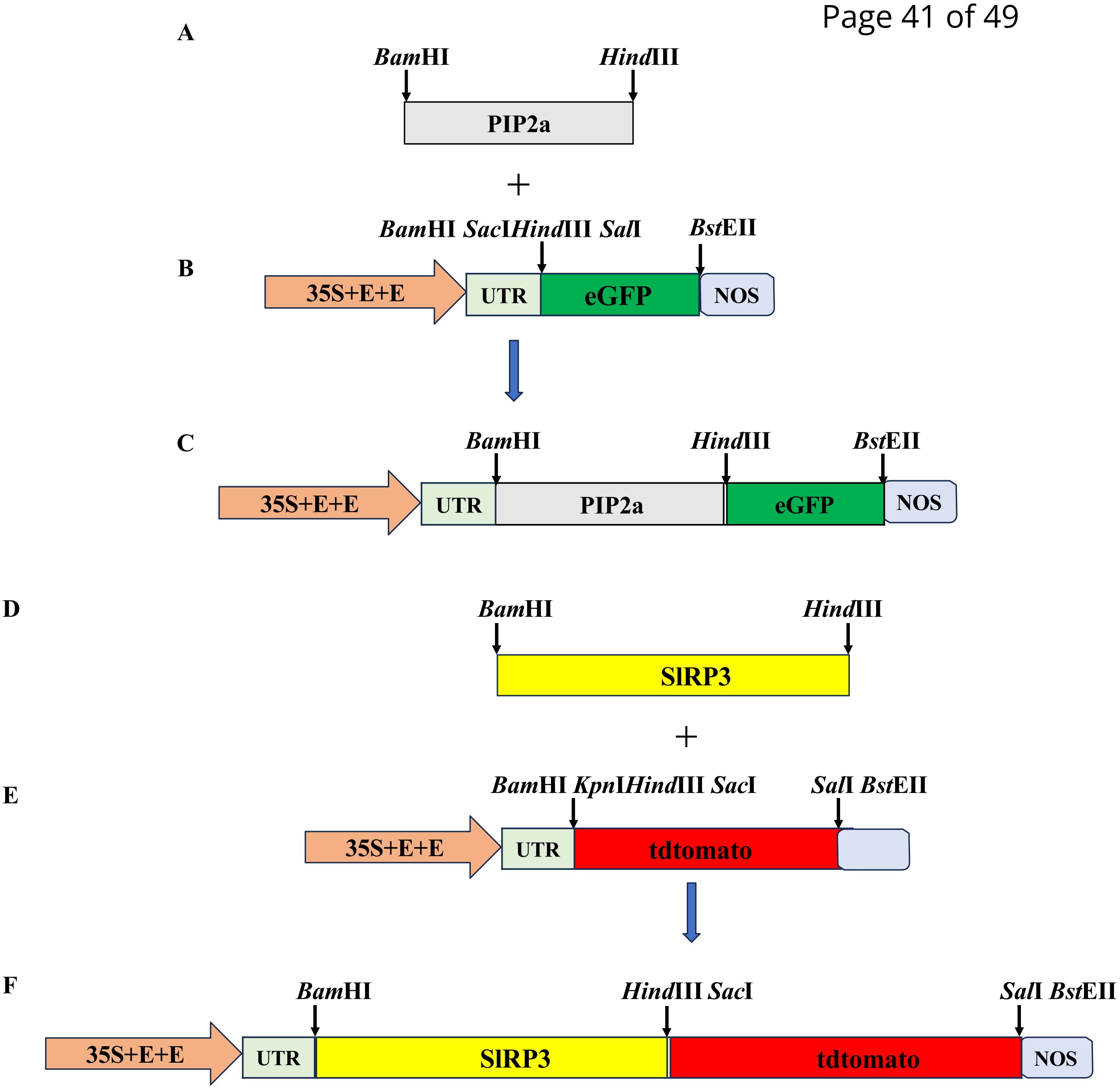


**Supplementary figure 1 :** Schematic representation of the cloning of PIP2a ORF into eGFP: pCAMBIA1304 (Plasma membrane marker) and SlRP3 into tdtomato: pCAMBIA1304 constructs. A and D: PCR amplification of PIP2a and SlRP3 without the stop codon with appropriate primers comprising BamHI and HindIII sites. B and E: Scheme representing vector backbone of eGFP :pCAMBIA1304 and tdtomato:pCAMBIA1304 respectively
C and F: Schematic representing positive clones of PIP2a: eGFP:pCAMBIA1304(Plasma membrane marker constructs) and SlRP3:tdtomato: pCAMBIA1304 respectively

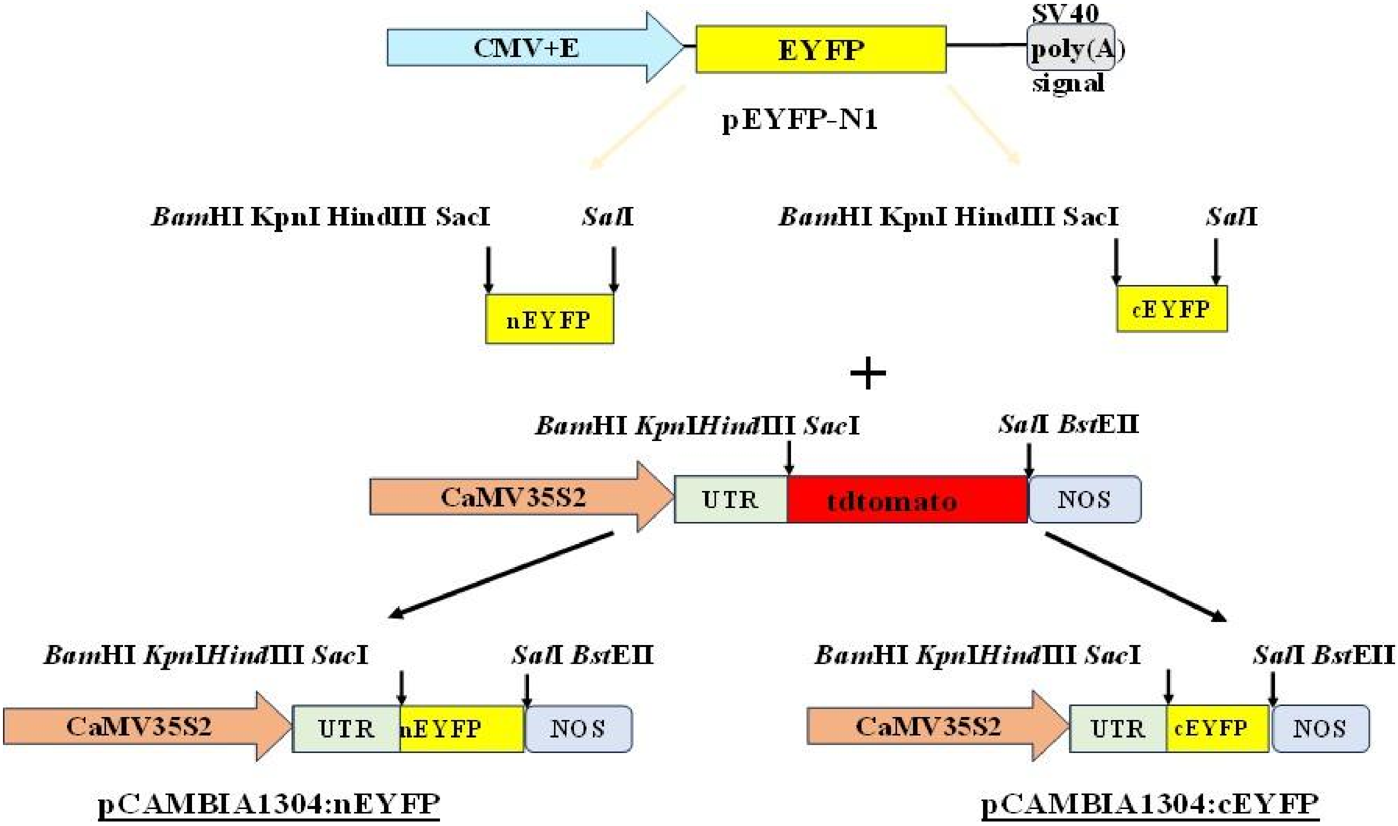


**Supplementary Figure 2:** Schematic representation of the insertion of nEYFP and cEYFP into pCAMBIA1304: tdtomato construct for generation of pCAMBIA1304: nEYFP and pCAMBIA1304: nEYFP constructs. The nEYFP and cEYFP sequence was PCR amplified from pEYFP-N1 vector with appropriate primers comprising of various restriction sites. The PCR products along with pCAMBIA1304: tdtomato vector was subjected to BamHI and SalI and thereafter ligated resulting in pCAMBIA1304: nEYFP and pCAMBIA1304: cEYFP vectors via cloning.

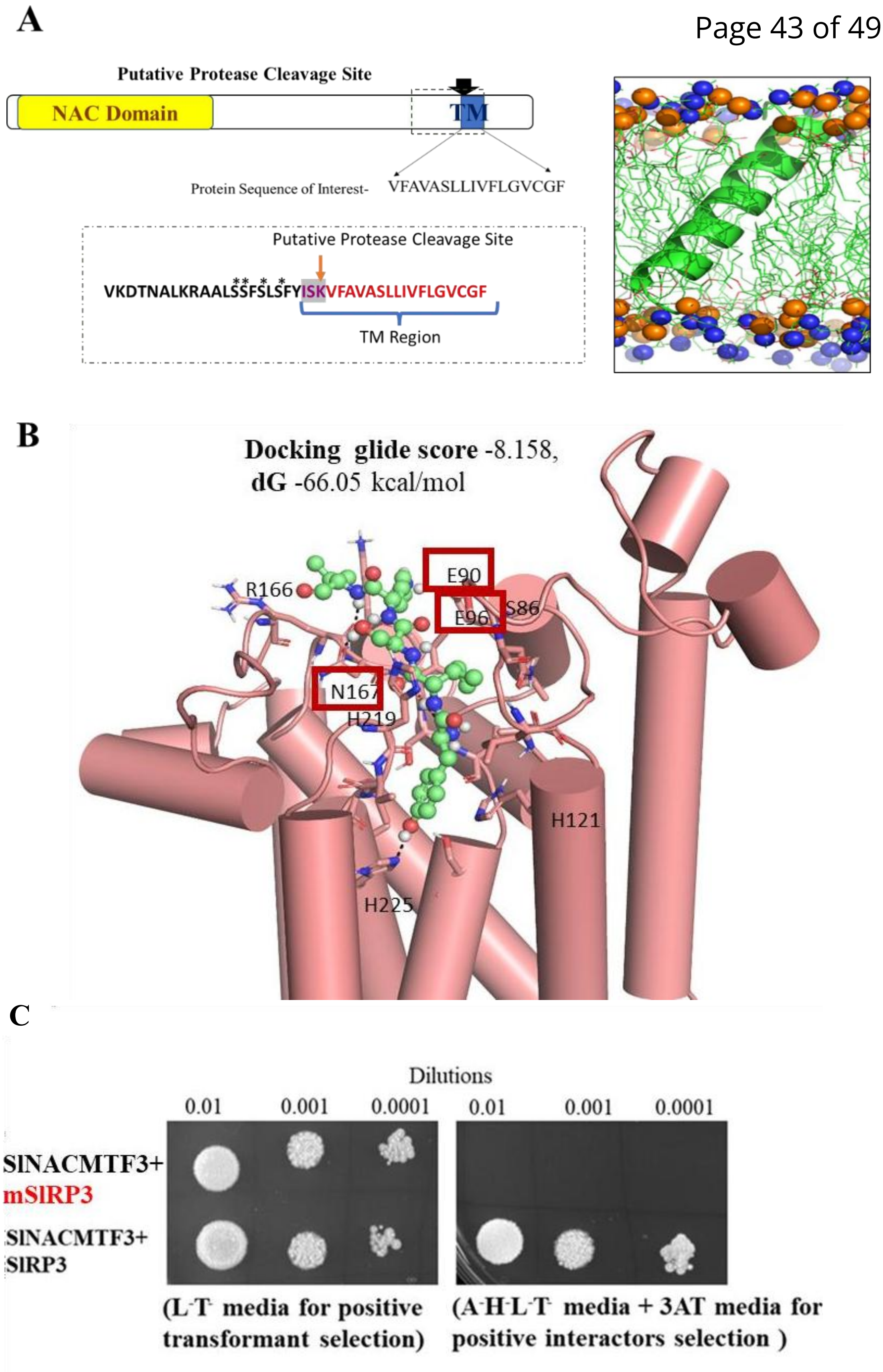


**Supplementary Figure 3:** Structural analysis of SlRP3 and TM domain of SlNACMTF3 A. In silico analysis of sequence of SlNACMTF3 and structure prediction of TM Region of SlNACMTF3.Sequence analysis reveals the presence of putative RP cleavage site upstream of TM region. The 17a.a after the Putative Protease Cleavage Site of SlNACMTF3 adopts a **helical** conformation spanning the membrane bilayer. The * represents presence of stretch of Serine amino acid near the putative protease cleavage site B. Docking model of SlRP3(protease) with pentapeptide (YISKV) of SlNACMTF3 (substrate)The ball and stick model represents pentapeptide residues of SlNACMTF3. Predicted amino acid residues of SlRP3 responsible for interaction with SlNACMTF3(substrate) are within red boxes. C. Mutated form of SlRP3 (mSlRP3:E90A+E96A+N167A) fails to interact with SlNACMTF3 which was validated using Split- Ubiquitin Membrane Based Yeast-Two Hybrid

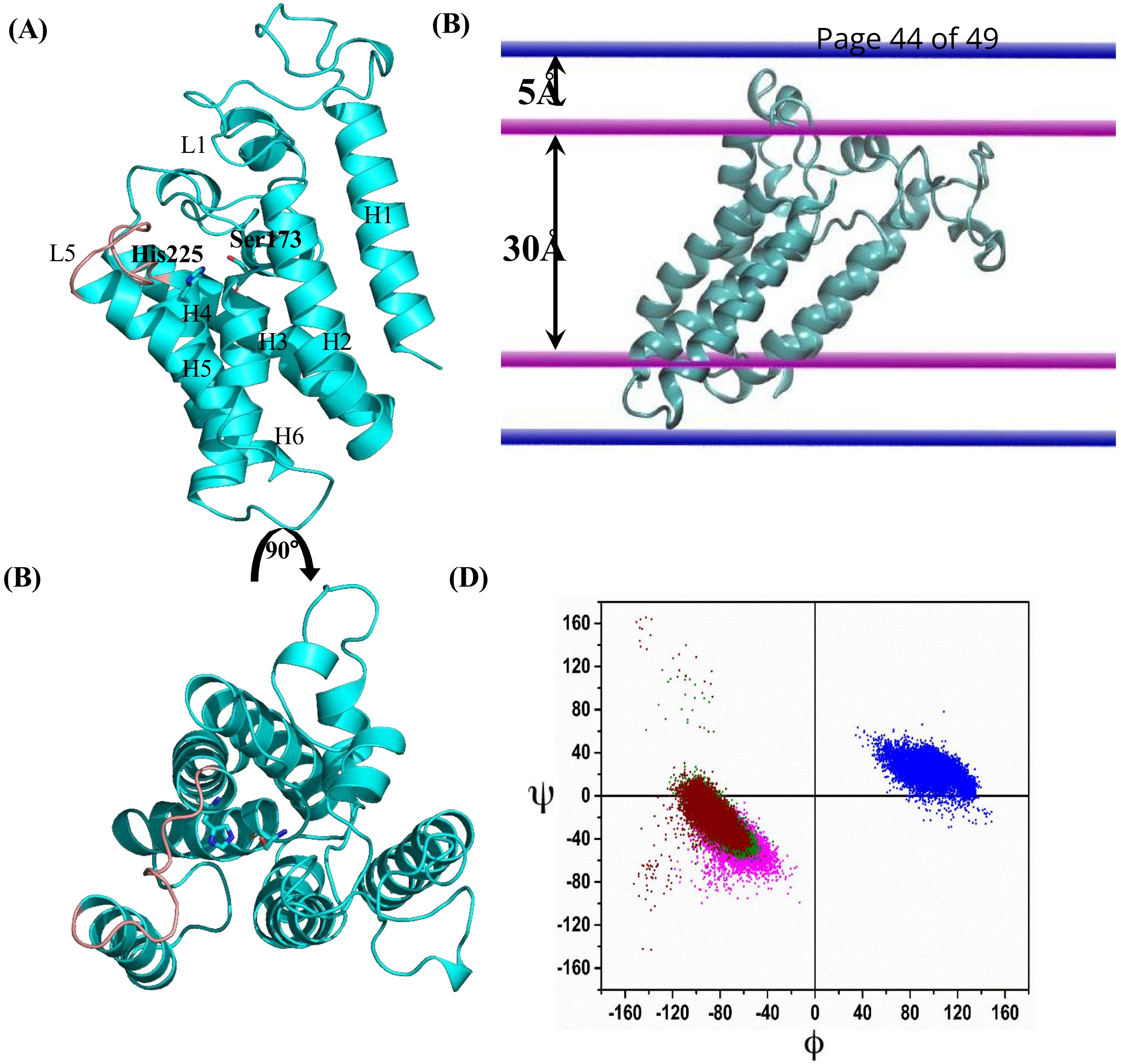


**Supplementary Figure 4 :** Structural characterization and validation of tomato heat- stress responsive Rhomboid Protease(SlRP3). A: Cartoon representation of structure of SlRP3 in (A) Side and (B) top view(present in main Figure also). The helices are marked from H1-H6. L1 and L5 represent loops 1 and 5; L5 is coloured in pink. Catalytic dyad residues S173-H225 are shown in sticks. (C). Alignment of SlRP3 in the membrane. The pairs of straight line indicate the boundary of membrane bilayer. (D) Ramachandran plot of residue 213 showing backbone Φ/Ψ sampling during MD simulation at different temperatures.

(A)

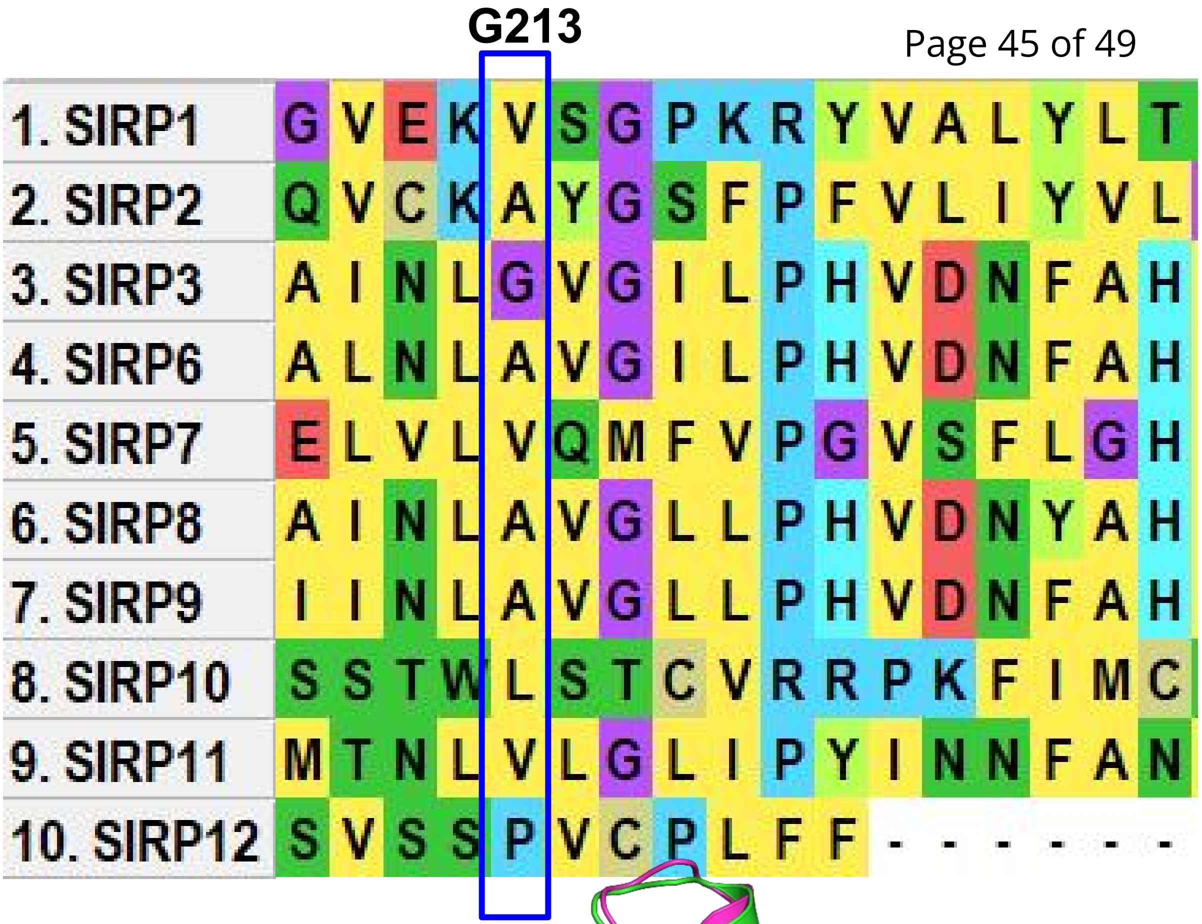


(B)

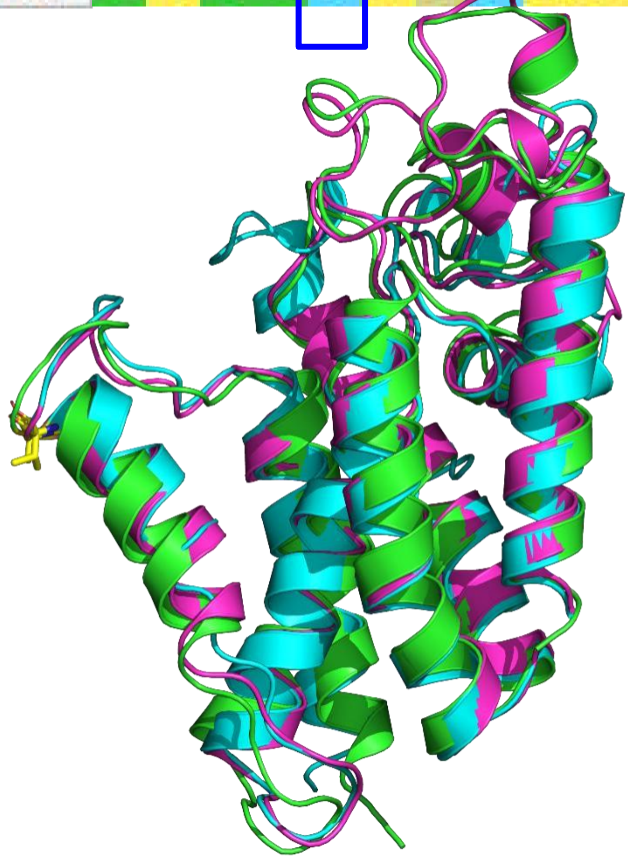

**Supplementary Figure 5**

(A) MSA of the L5 region showing unique position of G213 among all the characterised Rhomboid Proteases in tomato. (B) Cartoon representation of overlaid structures of all heat stress responsive SlRPs showing H5-L5 junction residues (yellow). Colour code of the proteins: green SlRP3, cyan SlRP7, and magenta SlRP9.

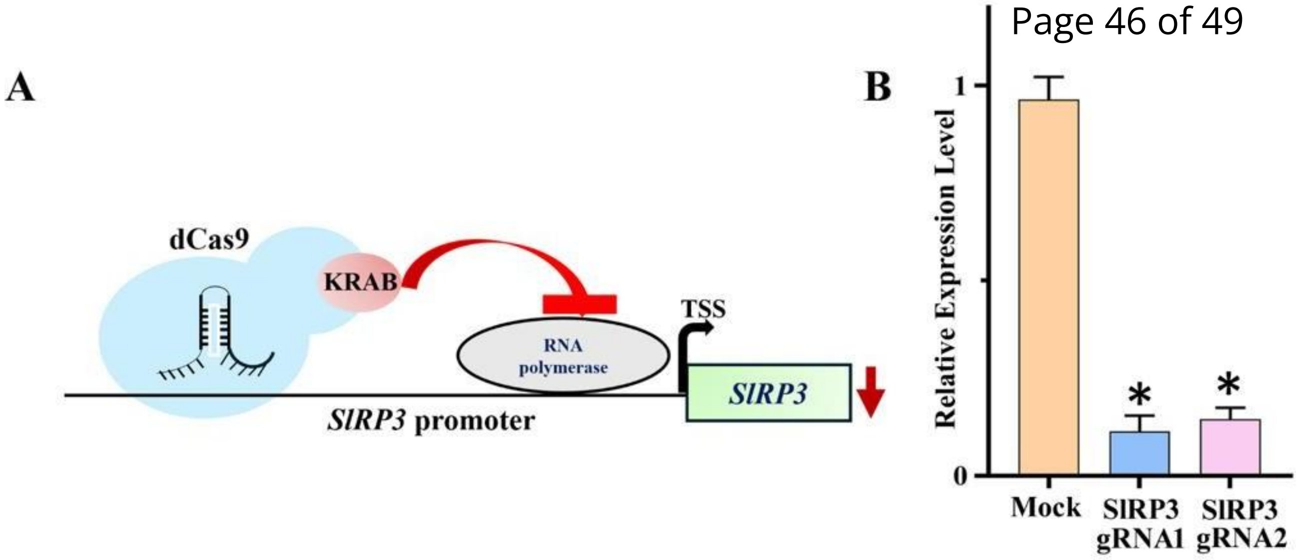


**Supplementary Figure 6:** CRISPR/dCas9(CRISPRi)-mediated transcriptional repression of SlRP3. (A) Schematic representation of the CRISPR interference (CRISPRi) strategy used to repress SlRP3 expression. A dCas9–KRAB repressor complex is targeted to the SlRP3 promoter, where it inhibits transcriptional initiation by interfering with RNA polymerase recruitment/activity. (B) Relative expression levels of SlRP3 in mock-treated plants and independent CRISPRi lines (SlRP3 gRNA1 and SlRP3 gRNA2), showing significant reduction of SlRP3 transcript abundance in both gRNA lines. Data are presented as mean ± SE. Asterisks indicate statistically significant differences compared with the mock control (*P < 0.05).

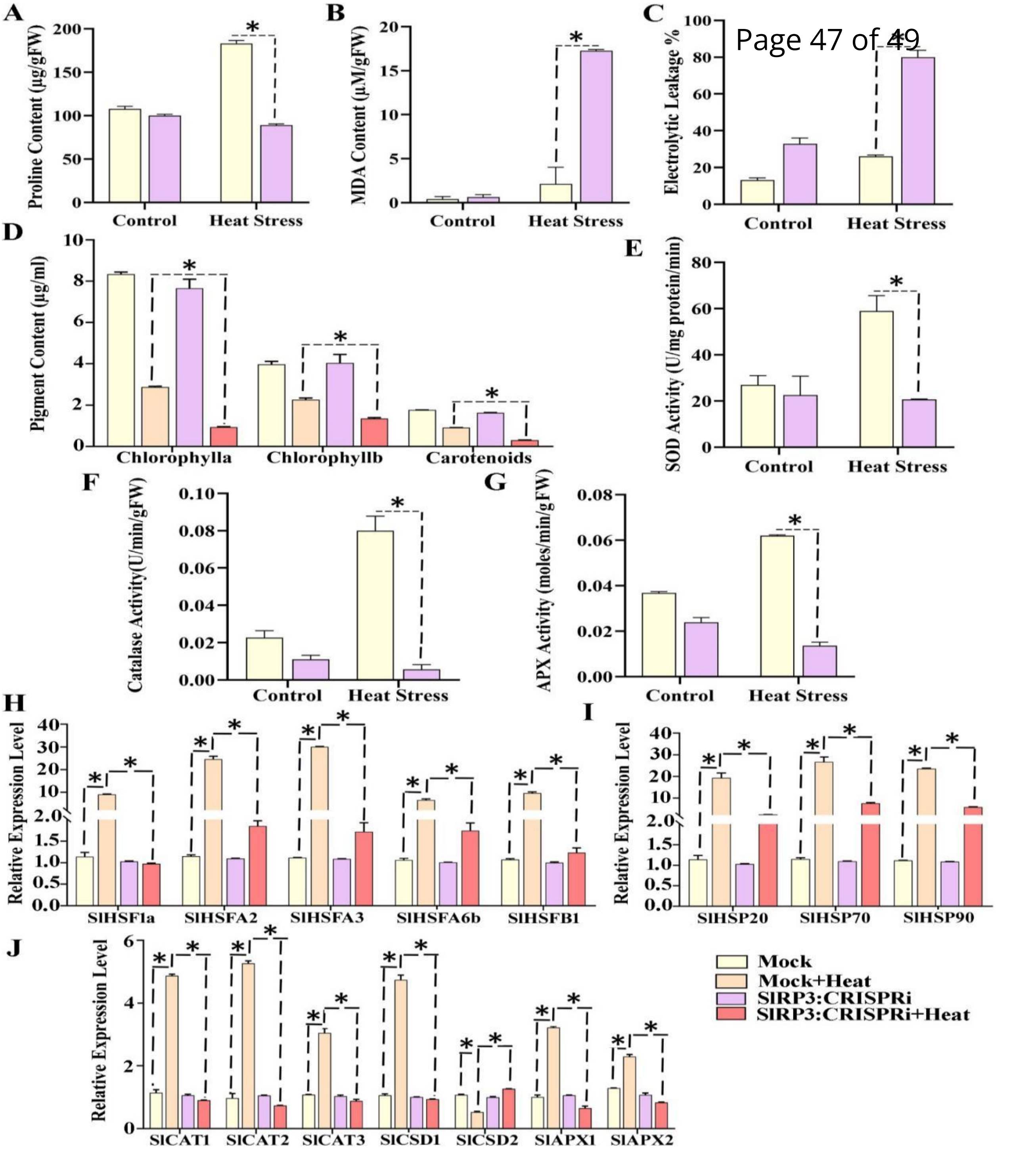


**Supplementary Figure 7** : SlRP3 is required for maintaining cellular homeostasis and antioxidant and heat-responsive pathways during heat stress. (A) Proline content in mock and SlRP3:CRISPRi plants under control and heat-stress conditions. (B) Malondialdehyde (MDA) accumulation, indicating heat-induced lipid peroxidation. (C) Electrolyte leakage as an indicator of heat-induced membrane damage. (D) Changes in photosynthetic pigments, including chlorophyll *a*, chlorophyll *b* and carotenoids. (E–G) Activities of antioxidant enzymes superoxide dismutase (SOD), catalase (CAT), and ascorbate peroxidase (APX), respectively, under control and heat-stress conditions. (H) Relative transcript levels of heat stress factor genes (SlHSF1a, SlHSFA2, SlHSFA3, SlHSFA6b and SlHSFB1). (I) Relative expression of heat shock protein genes (SlHSP20, SlHSP70 and SlHSP90). (J) Relative expression of antioxidant and stress-responsive genes, including SlCAT1, SlCAT2, SlCAT3, SlCSD1, SlCSD2, SlAPX1 and SlAPX2. The four treatments are represented by different bars: mock control, mock + heat, SlRP3:CRISPRi, and SlRP3:CRISPRi + heat, as indicated in the legend. Heat stress markedly alters osmoprotectant accumulation, membrane stability, photosynthetic pigments, antioxidant enzyme activities and expression of heat-responsive and ROS-scavenging genes in SlRP3:CRISPRi plants, indicating an important role of SlRP3 in the heat-stress response. Asterisks indicate statistically significant differences between the indicated groups. Data are presented as mean ± SD/SE (as applicable).

Supplementary Table 1: Biochemical and gene expression analyses data.

BIOCHEMICAL ASSAYS

| | Mock infiltrated vs Mock infiltrated heat | SlRP3:Knockdown C vs SlRP3:Knockdown H | Mock-infiltrated heat vs SlRP3:Knockdown H |
|---|---|---|---|
| **$H_2O_2$** | 5.53 ± 0.03 | 15.91 ± 0.04 | 2.76 ± 0.08 |
| **SOD activity** | 2.19 ± 0.61 | 0.92 ± 0.2 | 0.35±0.19 folds (p≤0.005) |
| **Catalase activity** | 3.47 ± 0.5 | 0.54 ± 0.78 | 0.1±0.87 |
| **APX activity** | 1.68 ± 0.001 | 0.58± 0.1 | 0.22 ±0.002 |
| **Electrolytic leakage** | 1.96 ± 0.29 | 2.42 ± 1.55 | 3.07±2.34 |
| **MDA content** | 5.19 ± 0.15 | 27.61± 0.17 | 8.304±0.11 |
| **Proline** | 1.70±0.98 | 0.89±1.24 | 0.48±3.87 |
| **Chlorophyll a content** | 0.34 ±0.1 | 0.121 ± 0.14 | 0.32±0.05 (p≤0.001) |
| **Chlorophyll b content** | 0.57 ± 0.18 | 0.33 ± 0.40 | 0.59±0.08 (p≤0.001) |
| **Carotenoid** | 0.51± 0.1 | 0.18± 0.23 | 0.33±0.01 (p≤0.001) |

**Antioxidant enzyme transcript regulation levels**

| | Mock infiltrated vs Mock infiltrated heat | SlRP3:Knockdown C vs SlRP3:Knockdown H | Mock-infiltrated heat vs SlRP3:Knockdown H |
|---|---|---|---|
| **SlCAT1** | 4.15±0.51 | 0.85± 0.33 | 0.185±0.04 |
| **SlCAT2** | 5.44± 0.46 | 0.69± 0.47 | 0.139±0.08 |
| **SlCAT3** | 2.81 ± 0.04 | 0.85 ±1.58 | 0.288±0.15 |
| **SlCSD1** | 4.46 ± 0.26 | 0.93 ± 0.2 | 0.197±0.15 |
| **SlCSD2** | 0.488 ± 1.25 | 1.26 ± 0.22 | 2.411+0.02 |
| **SlAPX1** | 3.21 ± 0.9 | 0.61 ±0.17 | 0.201±0.07 |
| **SlAPX2** | 1.78 ± 0.09 | 0.773 ± 0.25 | 0.361±0.06 |

**HSFs and HSPs transcript regulation levels**

| | | | |
|---|---|---|---|
| **SlHSFA1a** | 7.71 ± 0.11 | 0.95 ± 0.1 | 0.05±0.49 |
| **SlHSFA2** | 21.46 ± 1.13 | 1.67 ± 0.07 | 0.07±1.28 |
| **SlHSFA3** | 27.03 ± 0.09 | 1.60 ± 0.1 | 0.06±0.25 |
| **SlHSFA6b** | 6.3 ± 0.03 | 1.73 ± 0.2 | 0.25±0.53 |
| **SlHSFB1** | 9.11± 0.3 | 1.26± 0.1 | 0.13±0.50 |
| **SlHSP20** | 16.22 ± 1.40 | 2.23 ± 0.4 | 0.121 ± 0.38 |
| **SlHSP70** | 23.4 ±1.32 | 6.89 ± 0.18 | 0.28±0.32 |
| **SlHSP90** | 21.13 ± 0.67 | 5.33 ± 0.10 | 0.24±0.17 |

Summary statement: Heat-dependent conformational change of tomato rhomboid protease 3 facilitates SlNACMTF3 recognition within plasma membrane. The protease-mediated rapid activation leads to stress gene regulation for thermotolerance.
